# Control of a type III-Dv CRISPR‒Cas system by the transcription factor RpaB and interaction of its leader transcript with the DEAD-box RNA helicase CrhR

**DOI:** 10.1101/2023.12.07.570523

**Authors:** Raphael Bilger, Angela Migur, Alexander Wulf, Claudia Steglich, Henning Urlaub, Wolfgang R. Hess

**Affiliations:** University of Freiburg, Faculty of Biology, Genetics and Experimental Bioinformatics, Schänzlestr. 1, D-79104 Freiburg, Germany; Bioanalytics Research Group, Department of Clinical Chemistry, University Medical Centre, D-37075, Göttingen, Germany; Bioanalytical Mass Spectrometry Group, Max Planck Institute for Biophysical Chemistry, D-37077 Göttingen, Germany

**Keywords:** CRISPR-Cas systems, CrhR DEAD-box RNA helicase, CRISPR leader, cyanobacteria, regulation of CRISPR-Cas gene expression, RNA structure, transcription factor RpaB

## Abstract

CRISPR‒Cas systems in bacteria and archaea provide powerful defense against phages and other foreign genetic elements. The principles of CRISPR‒Cas activity are well understood, but less is known about how their expression is regulated. The cyanobacterium *Synechocystis* sp. PCC 6803 encodes three different CRISPR‒Cas systems. The expression of one of these, a type III-Dv system, responds to changes in environmental conditions, such as nitrogen starvation or varying light intensities. Here, we found that the promoter of the six-gene *cas* operon for the type III-Dv system is controlled by the light-and redox-responsive transcription factor RpaB. RpaB binds to an HLR1 motif located 53 to 70 nt upstream of the transcription start site, resulting in transcriptional activation at low light intensities. However, the strong promoter that drives transcription of the cognate repeat-spacer array is not controlled by RpaB. Instead, we found that the 125 nt leader transcript is bound by the redox-sensitive RNA helicase CrhR. Crosslinking coupled to mass spectrometry analysis revealed six residues involved in the CrhR-RNA interaction. Of these, L103, F104, H225, and C371 were predicted to be on the surface of a dimeric CrhR model, while C184 was not on the surface, and P443 could not be assigned to a structural element. These results showed that the expression of the CRISPR‒Cas system is linked to the redox status of the photosynthetic cyanobacterial cell at two different levels. While RpaB affects transcription, CrhR interacts with the leader transcript posttranscription. These results highlight the complex interplay between a CRISPR‒Cas system and its host cell.

## Introduction

CRISPR–Cas systems encode RNA-based adaptive and inheritable immune systems in many archaea and bacteria^1,2^; these systems are highly diverse and were classified into two classes, six types and 33 subtypes^3^; however, new subtypes are still being discovered. Type III CRISPR‒Cas systems, characterized by the presence of the signature gene *cas10*, exist in 34% and 25% of archaeal and bacterial genomes that encode CRISPR‒Cas loci, respectively^4^. Type III systems are further classified into five subtypes, A to E^3,5,6^. Although detailed insights have been obtained regarding the molecular mechanisms and peculiarities of the different types of CRISPR‒Cas systems, knowledge about how their expression is regulated has remained incomplete.

In the subtype I-E system of *E. coli*, regulation by transcription factors has been demonstrated. The DNA-binding protein HNS (Histone-like Nucleoid Structuring Protein) acts as a repressor by inhibiting the expression of crRNA and *cas* genes^7^. As an antagonist of HNS, LeuO activates the expression of *cas* genes, thereby enhancing resistance against invading DNA^8^. Finally, a signaling cascade involving the BaeSR two-component regulatory system, which senses envelope stress (e.g., phage attack) via the membrane-localized kinase BaeS, was identified. Once activated, BaeS phosphorylates the cytoplasmic transcription factor BaeR^9^, which, among other genes, activates the expression of *cas* genes^10^.

In the thermophilic archaeon *Sulfolobus islandicus*, the expression of the type I-A CRISPR locus is regulated by Csa3a and Csa3b, two transcriptional regulators containing CARF and HTH domains. While Csa3a activates the expression of adaptation genes and the CRISPR array^11^, interference genes are repressed by Csa3b. Repression is achieved in the absence of viral infection by cobinding of the cascade complex^12^. In Serratia, the LysR-type transcriptional regulator PigU co-ordinately controls the expression of a type III-A and a type I-F system^13^. Further regulatory mechanisms have been described for CRISPR-Cas systems in *Pectobacterium atrosepticum*^14^ and *Pseudomonas aeruginosa*^15,16^.

Cyanobacteria are the only prokaryotes whose physiology is based on oxygenic photosynthesis, making them immensely important primary producers. Field studies have shown that both cyanobacterial cell counts and the number of coinfecting bacteriophages (cyanophages) can be very high, with up to 50% of all cyanobacteria estimated to be infected at any one time^17^, and likely affect cyanobacterial biogeography and biogeochemistry at the scale of oceanic subregions^18^. Accordingly, active defense mechanisms can be expected in cyanobacteria.

The unicellular cyanobacterium *Synechocystis* sp. PCC 6803 (from here: *Synechocystis* 6803) is a model for the CRISPR biology of cyanobacteria. It possesses three separate and complete CRISPR‒Cas systems, a type I-D (CRISPR1), III-Dv (CRISPR2), and III-Bv (CRISPR3) system, which are highly expressed under a variety of conditions and active in interference assays^3,19–25^. The CRISPR2 system is of particular interest because it has recently been suggested to function as a protein-assisted ribozyme^25^.

Each of the three CRISPR‒Cas loci in *Synechocystis* 6803 is associated with one gene that has been suggested to be a regulator (genes *sll7009*, *sll7062* and *sll7078*^19^). Indeed, deletion of *sll7009*, which encodes a putative WYL domain protein, led to increased accumulation of crRNAs in the CRISPR1 system but did not change the crRNA levels in the other two systems^26^. This result was consistent with the observation that CARF and WYL domain regulatory proteins are widely distributed ligand-binding specific regulators of CRISPR‒Cas systems^27^. Sll7062 differs from the other two possible regulators by the presence of an N-terminal CARF7 family domain fused to a RelE RNase domain, a setup characteristic of Csm6 proteins. Csm6 proteins are not transcription factors but rather CRISPR-associated RNases that are activated by cyclic oligoadenylate (cOA)-mediated signaling^28^. Accordingly, Sll7062 was renamed SyCsm6 when its activity was tested upon production as a recombinant protein, together with the CARF-HEPN domain protein SyCsx1 (Slr7061)^29^. Therefore, the CRISPR2 system lacks an obvious candidate regulatory gene in its vicinity.

However, when we characterized the regulon controlled by the transcription factor RpaB, we noted the possible involvement of a host genome-encoded factor in CRISPR2 regulation^30^. RpaB (‘‘regulator of phycobilisome association B”, Slr0947) is an OmpR-type transcription factor that is predicted to control more than 150 promoters by binding to the HLR1 (‘‘high light regulatory 1’’) motif, a pair of imperfect 8-nt long direct repeats (G/T)TTACA(T/A) (T/A) separated by two random nucleotides. RpaB mediates transcriptional activation when the HLR1 motif is located 45 to 66 nt upstream of the transcription start site (TSS), whereas all other locations mediate repression^30^. The results showed that RpaB is a transcription factor of central importance for light-and redox-dependent remodeling of the photosynthetic apparatus and many associated pathways. Surprisingly, there was also a predicted binding site in the promoter that drives the transcription of the *cas* gene operon of CRISPR2, the III-D system in *Synechocystis* 6803, but this was not investigated further.

Another protein with a central role in light-and redox-dependent responses in *Synechocystis* 6803 is the cyanobacterial RNA helicase Redox (CrhR)^31^. CrhR (Slr0083) is the single DEAD-box RNA helicase in *Synechocystis* 6803 that is capable of altering RNA secondary structures by catalyzing double-stranded RNA unwinding as well as annealing^32^. The molecular effects of *crhR* deletion or inactivation have been studied at the transcriptome^33,34^ and proteome levels^35^, and several attempts have been made to identify the RNA targets of CrhR directly^36^.

Here, we applied a further approach to pull down RNA that interacts with CrhR, which is expressed as a recombinant protein, and found that the transcribed leader of the type III-Dv CRISPR‒Cas system was copurified. Therefore, we investigated the possible regulatory impact of the host genome-encoded transcription factor RpaB on the expression of the CRISPR2 system and described and validated the interaction of CrhR with the leader transcript of the repeat-spacer array of the same system.

## Results

### The expression of the type III-Dv CRISPR2 system in *Synechocystis* 6803 is affected by environmental conditions

In our previous analysis of the distribution of putative HLR1 binding sites for the transcription factor RpaB in *Synechocystis* 6803, one site was predicted in the CRISPR2 *cas* gene promoter; however, this site has not been studied further^30^. This promoter drives the transcription of six genes, *sll7067* to *sll7062*, into a single transcriptional unit (TU)^37^. Therefore, these six genes constitute an operon. These genes encode Cas10, a Cas7-Cas5-Cas11 fusion, Cas7-2x, Csx19, Cas7 with an insertion, and the SyCsm6 protein (**Figure 1A**). Technically, two TUs, TU7058 and TU7063, were defined for the CRISPR2 *cas10* promoter because they contain two TSSs (at positions 62704 and 62807 on the reverse strand)^37^. Our previous genome-wide mapping of TSSs using differential RNA-Seq indicated the regulated expression of this operon. High numbers of reads were found for TU7058 under most of the tested growth conditions, but relatively lower numbers were recorded after the cultures were transferred to high light (470 μmol photons m^-2^ s^-1^ for 30 min), and no reads were detected at all if the cultures were incubated in the dark for 12 h^37^.

**Figure 1.**
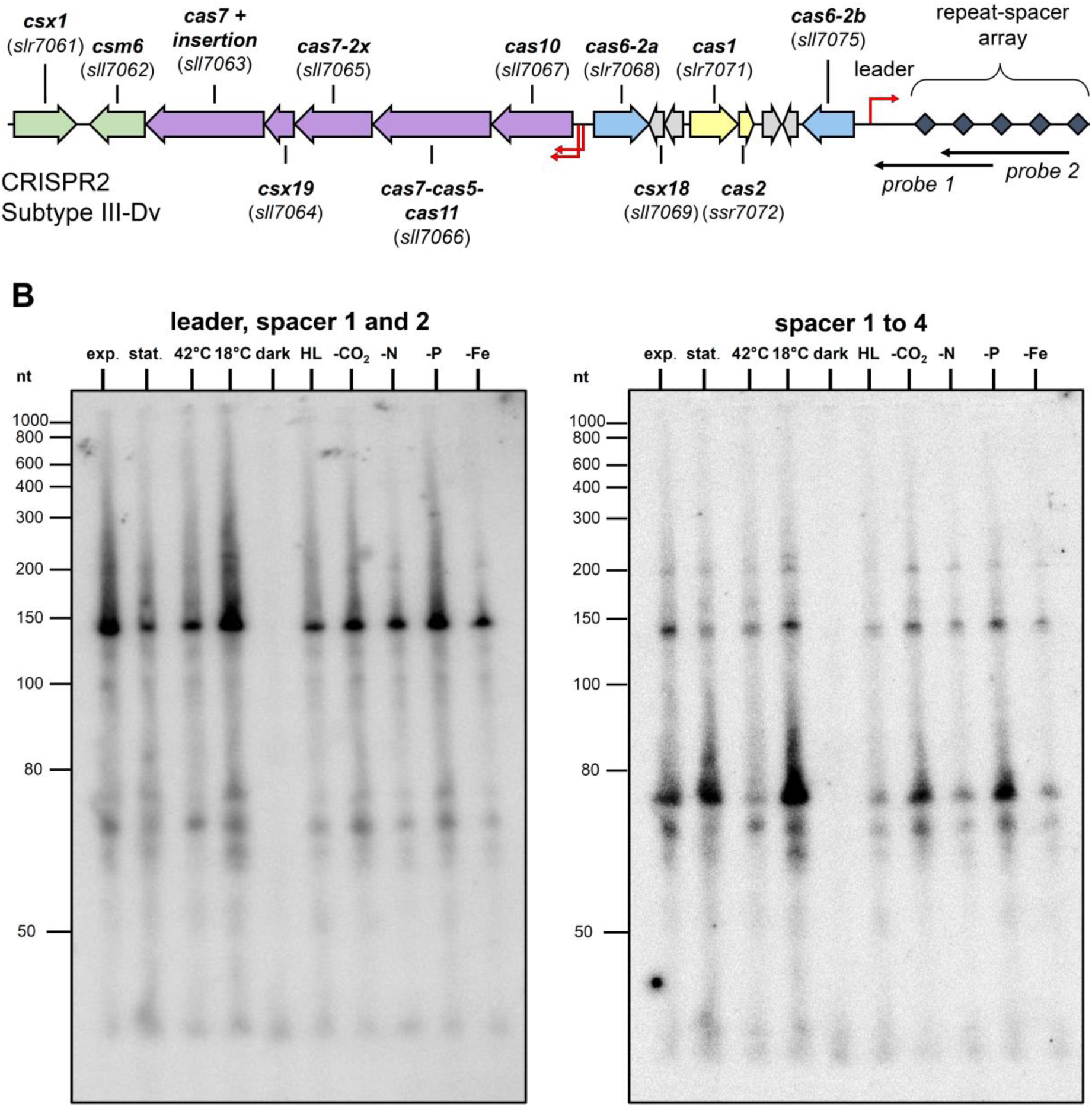
Organization of the type III-Dv (CRISPR2) locus in *Synechocystis* 6803 and the influence of environmental conditions. **A.** The type III-Dv CRISPR‒Cas system is located on the pSYSA plasmid. Several *cas* genes are located upstream of the CRISPR array, which consists of a 125 nt long leader and 56 spacers 34-46 nt in length interspaced by 37 nt long repeats (gray squares). Arrows in yellow indicate *cas* genes encoding proteins for the adaptation module; blue, *cas6* genes; and purple, genes encoding the effector complex. Accessory genes are indicated by green arrows, and genes encoding hypothetical proteins are shown in light gray. The transcriptional start sites of the CRISPR array and the effector complex operon are marked by bent red arrows, and the locations of the antisense RNA probes used for Northern hybridization are indicated by straight narrow arrows in black. **B.** Influence of different environmental conditions on the accumulation of leader and crRNAs, including spacers 1 and 2 (left panel) and spacers 1 to 4 (right panel). For Northern hybridization, ^32^P-labeled transcript probes, as indicated in panel (A), were used after separation of 12 µg of RNA each isolated from cultures grown under 10 different conditions on a 10% urea‒polyacrylamide gel. Exp. (exponential phase), stat. (stationary phase), 42 °C (heat stress), 18 °C (cold stress), dark (darkness), HL, high light (470 μmol photons m^-2^ s^-1^ for 30 min), -CO_2_ (limitation in inorganic carbon supply), -N (nitrogen limitation), -P (phosphorus limitation) and -Fe (iron limitation). The membrane and 5S rRNA hybridization to control equal loading are shown in Figure 2A of publication^76^.

The respective repeat spacer array is transcribed on the forward strand, starting from a single TSS approximately 6 kb away from the *cas* gene operon (**Figure 1A**). To explore the possible differential accumulation of leader and CRISPR RNAs (crRNAs), total RNA samples obtained from cultures grown under the same ten conditions as those previously used for differential RNA-Seq were analyzed via Northern hybridization. We used two probes, complementary to the CRISPR leader RNA and the first two spacers and repeats or to spacers 1 to 4. The first probe produced a major signal of approximately 150 nt (**Figure 1B**, left panel), which matches the length of the leader (125 nt;^19^) plus the length of the cleavage site within the first repeat (27 nt;^20^), and two weaker signals matching the lengths of a repeat-spacer unit of ∼72 nt and the final processed spacer 1 of 44 nt. The second probe detected the same ∼150 nt precursor transcript due to overlap in the repeat but revealed the strongest signals for repeat spacer units 2 and 3, which are somewhat longer (∼75 to 77 nt) than other units. Their accumulation was highly dependent on the conditions. The strongest signals were obtained with the samples from cultures exposed to cold stress, stationary phase, N, and C starvation, whereas the signals were weaker in samples from cultures exposed to heat shock or high light and were not detected in samples from cultures incubated in the dark for 12 h (**Figure 1B**, right panel). These results matched the differential transcript accumulation observed for the CRISPR2 *cas10* operon via differential RNA-Seq.

### Transcriptional regulation of the CRISPR2 *cas 10* promoter

The observed differential accumulation of *cas* gene operon-mRNAs and crRNAs may be due to differential transcription, posttranscriptional regulation, or both. The prediction of a putative HLR1 motif in the CRISPR2 *cas10* promoter indicated possible transcriptional regulation. This HLR1 motif is located -70 nt to -53 nt from the TSS of TU7063 and -172 to -155 nt from the TSS of TU7058. To examine its possible relevance, we cloned the 5’UTR of the CRISPR2 *cas* gene and the promoter region (+122 to -203 with regard to the TSS of TU7058), which included the HLR1 motif (native promoter, P_nat_) upstream of the *luxAB* reporter gene in the vector pILA^38^. As a control, we mutated the HLR1 motif by substituting four nucleotides with guanosines (mutated promoter, P_mut_). Initial P_nat_ activity was measured under low-light conditions. Promoter activity was measured again after a 4-hour incubation under high light, where we observed a decrease in the activity to the level of the no-promoter control (P_less_). After transfer back to low light, P_nat_ activity increased significantly over time, reaching an approximately tenfold increase in luminescence after 120 min (**Figure 2A**). In contrast, the P_mut_ promoter harboring the mutated HLR1 motif exhibited a basal level of bioluminescence, similar to that of the control strain harboring promoterless (P_less_) *luxAB* genes, even after the P_mut_ strain was transferred back to low light. This finding indicates the importance of mutated nucleotides in the recognition and binding of RpaB to the promoter. When we exposed the cells after the initial 4 h continuously to high light (**Figure 2B**) or added the electron transfer inhibitor DCMU (**Figure 2C**), bioluminescence remained at a basal level with P_nat_, P_mut_, and P_less_ for the duration of the experiment. This effect might be specifically related to RpaB and a change in redox status, as the P*_syr9_* promoter used for control had modest activity under high light conditions and was not negatively influenced by the added DCMU.

**Figure 2.**
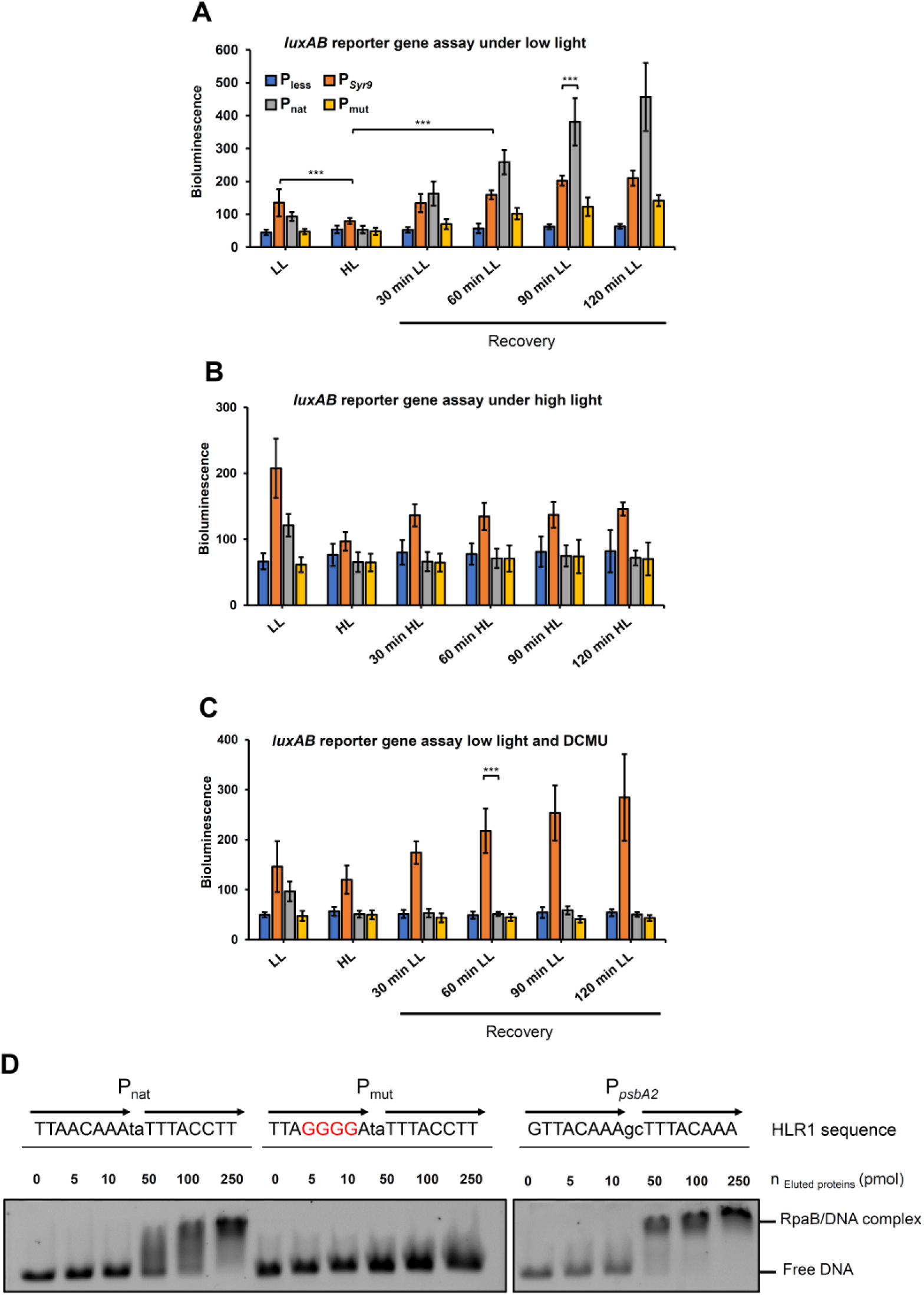
RpaB control of the CRISPR2 *cas10* gene promoter. **A.** Activity of the CRISPR2 *cas10* promoter under continuous low light after exposure to high light. **B.** Activity of the CRISPR2 *cas10* promoter after the shift to high light. **C.** Activity of the CRISPR2 *cas10* promoter under continuous low light in the presence of DCMU after exposure to high light. *Synechocystis* strains were transformed with pILA constructs with a promoterless *luxAB* (P_less_), the sRNA Syr9 promoter (P*_syr9_*), the wild type (P_wt_) and the modified *cas10* promoter mutated in its HLR1 site (P_mut_). The bioluminescence of 100 µL culture aliquots was measured using a Victor^3^ multiplate reader. The data are presented as the means ± SDs from three independent experiments. **D.** Electrophoretic mobility shift assays (EMSAs) were used to test the binding of purified His-RpaB (**Figure S1**) to CRISPR2 *cas10* promoter fragments containing the native (P_nat_) or mutated HLR1 sequence (P_mut_). The HLR1-containing *psbA2* promoter fragment (P*_psbA2_*) was used as a positive control. Substituted bases in the P_mut_ fragment are highlighted in red. Then, 0.5 pmol Cy3-labeled DNA fragments 80 nt in length were incubated for 15 min in the dark with His-RpaB at the indicated concentrations. The samples were separated on 0.5x TBE and 3% agarose gels. The arrows represent imperfect repeats at the HLR1 site. HL: high light; LL: low light.

These results showed that the CRISPR2 *cas10* promoter is regulated by a redox-dependent mechanism involving the HLR1 motif. Furthermore, these results are consistent with the prediction that RpaB positively regulates this promoter because it is known to dissociate from its HLR1 binding motif under high light^39^.

### RpaB binds to the native CRISPR2 *cas10* promoter but not to the mutated HLR1 site

We then validated the prediction that RpaB regulates the transcription of the CRISPR2 effector complex. Therefore, RpaB from *Synechocystis* 6803, fused to a C-terminal 6×histidine tag, was expressed in *E. coli* DE3^39^ and purified using nickel chromatography (**Figure S1**). For the electrophoretic mobility shift assay (EMSA), increasing amounts of purified RpaB were incubated with 0.5 pmol of Cy3-labeled DNA probes harboring either the wild-type or the mutated HLR1 motif (P_nat_ and P_mut_). As a positive control, we used the *psbA2* promoter P*_psbA2_*, which was previously characterized and shown to contain a functional HLR1 motif^40^.

For the P_nat_ and P*_psbA2_* fragments, a band shift was observed with 50 pmol of recombinant 6×His-RpaB. For the P_mut_ fragment with four substituted bases within the HLR1 motif, the highest amount of added 6×His-RpaB (250 pmol) was not sufficient to induce a band shift (**Figure 2D**).

Taken together, these results strongly suggested that the redox-dependent transcription factor RpaB positively regulates the transcription of the CRISPR2 effector complex under low light conditions by binding to the HLR1 site. This finding implied that the expression of the CRISPR2 *cas10* complex is activated under low-light conditions by RpaB and deactivated under high light conditions when RpaB binding is lost. We wondered what this would mean to the accumulation of crRNAs. Moreover, upon acclimation to high light, RpaB regains DNA-binding activity^30^. Therefore, we performed another experiment in which we extended the time at high light to 6 h, transferred the cells to nitrogen starvation conditions, added the electron transport inhibitors DCMU or DBMIB to the cultures, and analyzed the accumulation of the CRISPR2 leader and crRNAs. Northern hybridization against spacers 1-4 produced several bands ranging from approximately 250 nt (pre-crRNA) to 72 nt (**Figure 3A**), which corresponded to a single-unit crRNA precursor^19^. At approximately 150 nt, we observed a double band corresponding to the partially processed pre-crRNAs, as also found in **Figure 1B**. Because the spacers differ in sequence and length, the intermediate cleavage products are slightly different in size. Contrary to the results in **Figure 1B**, we observed no decrease but a slight increase in the accumulation of pre-crRNA after six hours of exposure to high light, which indicated that the cells had acclimated to the new environmental conditions (**Figure 3A**). In nitrogen-depleted medium, we observed a decrease in pre-crRNA accumulation after six hours, consistent with the findings in **Figure 1B** and previous transcriptome analysis results^37^. To test the impact of changes in redox conditions on pre-crRNA accumulation, we added the photosynthesis inhibitor DCMU (3-(3,4-dichlorophenyl)-1,1-dimethylurea) or the cytochrome b_6_f complex inhibitor DBMIB (2,5-dibromo-3methyl-6-isopropylbenzoquinone) to our cultures. Here, we observed a weaker accumulation of the lower double bands at 150 nt. Furthermore, in the presence of DBMIB, mature crRNAs (< 80 nt) vanished almost completely, which is consistent with the overall decrease in spacer transcript accumulation in the presence of the inhibitors. To test whether spacer and leader accumulation differed, we hybridized the same membrane against the leader transcript (**Figure 3B**). The signal ran at approximately 150 nt, matching the previously estimated length of 125 nt for the leader transcript plus the length of the first repeat up to the first Cas6 cleavage site of 29 nt^19^. The accumulation of the leader was similar to that of spacer repeats under high light and nitrogen depletion conditions (**Figure 3A**). The addition of DCMU greatly reduced the accumulation of the leader compared to the standard (low light) conditions, and the addition of DBMIB resulted in the loss of the leader transcript signal. The observed effects of DCMU and DBMIB could be explained by a general inhibitory effect on RNA synthesis. To test this possibility, we hybridized a probe for *atpT* mRNA, which was previously found to be strongly induced by the addition of DCMU or DBMIB^41,42^. Both inhibitors upregulated the accumulation of *atpT* mRNA, demonstrating that transcription was not inhibited globally (**Figure 3C**).

**Figure 3.**
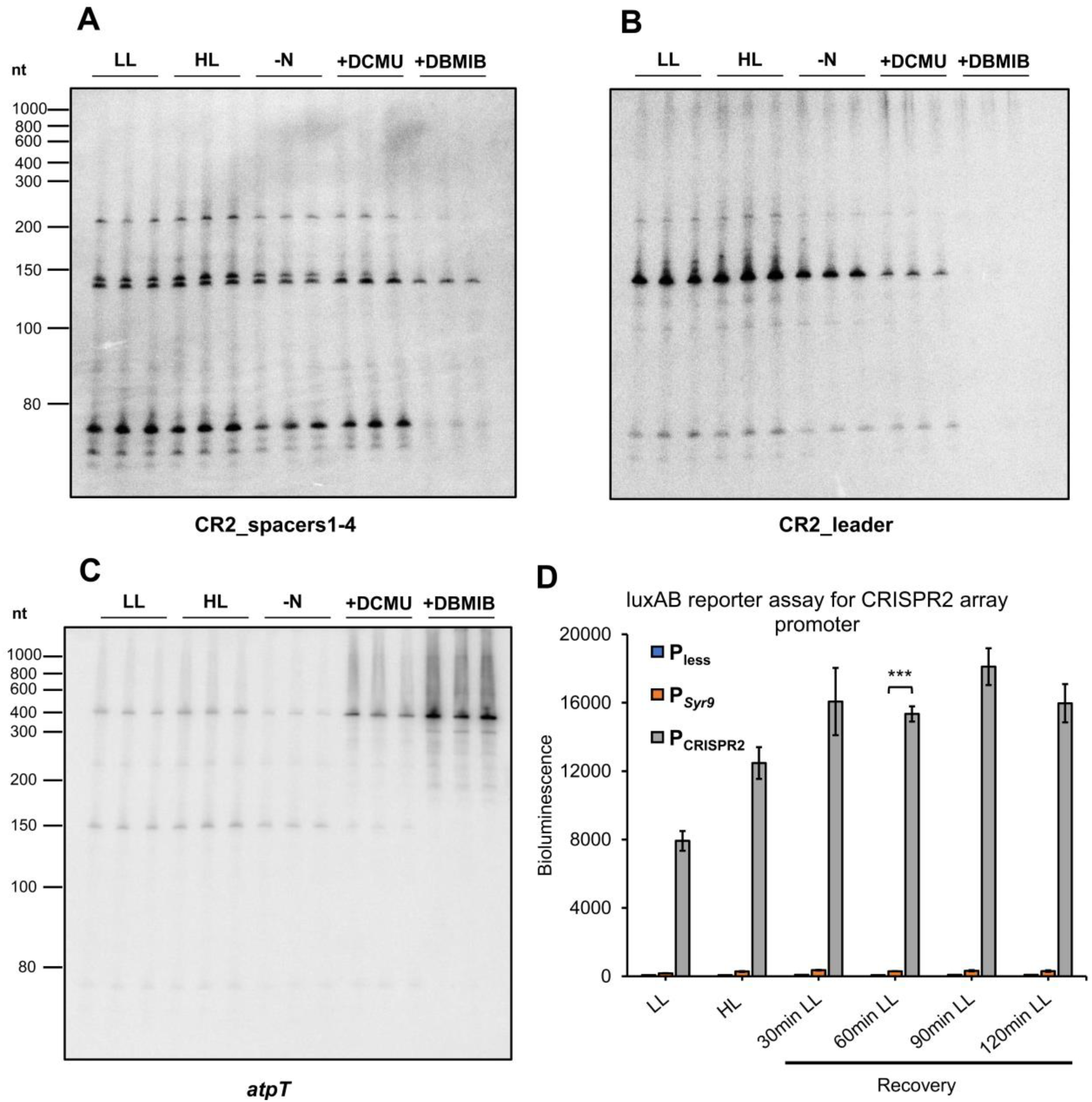
Effect of the addition of electron transport inhibitors on CRISPR leader accumulation. *Synechocystis* 6803 wild type was cultivated under low light (LL), high light (HL), and nitrogen-depleted BG11 media (-N) and in the presence of the electron transport inhibitors DCMU or DBMIB (50 µM each). The cells were harvested after 6 h of incubation. Total RNA was extracted, and 10 µg per lane was loaded onto an 8 M urea-10% PAA gel. Single-stranded RNA probes were hybridized against spacers 1-4 of CRISPR2 (A), the CRISPR2 leader (B), and *atpT* mRNA (C). The relative amounts of CRISPR2 leader transcripts were normalized to the 5S rRNA intensity and quantified. **D.** Activity of the CRISPR2 array promoter under continuous low-light conditions or after exposure to high light.

These results suggest that the stability of the CRISPR2 leader transcript is linked to the redox status of the plastoquinone pool.

### The CRISPR2 array promoter is highly active

We observed a decreased accumulation of the spacer-repeat and leader transcripts when the cells were exposed to high light (470 μmol photons m^-2^ s^-1^) for 30 min (**Figure 1B**). To determine whether the promoter itself might be regulated, similar to the promoter of the *cas* gene operon, we cloned the region -1 nt to -100 nt from its TSS, fused it to a synthetic ribosome binding site in the pILA vector, and integrated the construct in *Synechocystis 6803* as for the CRISPR2 *cas10* promoter constructs. **Figure 3D** shows the *luxAB* reporter assay with the CRISPR2 array promoter. The cultures were exposed to high light for 30 min before being returned to low light to avoid acclimation to the increased light intensity. The measured bioluminescence was extremely high, reaching 18,000 units, the highest measured activity in a comparison of five different promoters (**Figure S2**).

Moreover, we did not observe a decrease in the bioluminescence signal after exposing the cultures to high light for 30 min but rather a further increase in bioluminescence. These results suggested that the CRISPR2 array promoter is not influenced by environmental light conditions and that the observed changes in leader and crRNA transcript levels were caused by another mechanism.

### The CRISPR2 leader RNA interacts with CrhR

Because we found no evidence for RpaB controlling the crRNA promoter, we considered preliminary results that indicated the involvement of the DEAD-box RNA helicase CrhR as another possible factor. CrhR mediates light-and redox-dependent responses in *Synechocystis* 6803^31^. We used CrhR produced as a recombinant protein in *E. coli*. Two *E. coli* strains expressing recombinant His-tagged native CrhR or CrhR_K57A_ with enhanced RNA binding due to the K57A substitution within the ATP-binding motif were utilized. The ∼55 kDa proteins corresponding to His-tagged CrhR and CrhR_K57A_ were detected three hours after induction with 1 mM IPTG, purified via HiTrap Talon crude column (Cytiva) chromatography, and eluted with a step gradient of imidazole concentrations (**Figure S3**).

The recombinant proteins were incubated with *Synechocystis* 6803 total RNA and subjected to coimmunoprecipitation (co-IP), after which three cDNA libraries were prepared from the bound RNA, representing RNA interacting with recombinant CrhR, recombinant CrhR_K57A_, and total RNA as a background control. The experiment was performed in biological duplicates. The total numbers of reads obtained from the single-end Illumina sequencing are listed in **Table S1**. The reads were trimmed, and the adapter contaminants were filtered out with cutadapt and subsequently mapped to the *Synechocystis* 6803 chromosome and plasmids using Bowtie2^43^. Using the PEAKachu peak caller^44^, 39 peaks were identified in the CrhR library (**Figure 4**, **Table 1**), and 41 peaks were called with the RNA obtained from CrhR_K57A_ (**Figure 4**, **Table 2**), which met a log_2_FC ≥1 and adjusted *p* value ≤0.05. The peaks mapped to positions on the chromosome and the plasmids pSYSA, pSYSM and pSYSX. Of these, 24 peaks were shared between the two proteins including the CRISPR2 leader RNA (**Figure 4**). Both the RNA helicase CrhR and the CrhR_K57A_ mutant strongly interacted with their own mRNAs, consistent with previous results on its autoregulatory features^45^. In addition to those of the leader, several crRNAs of the CRISPR2 array were also enriched in the CrhR co-IP (**Table 1**).

**Figure 4.**
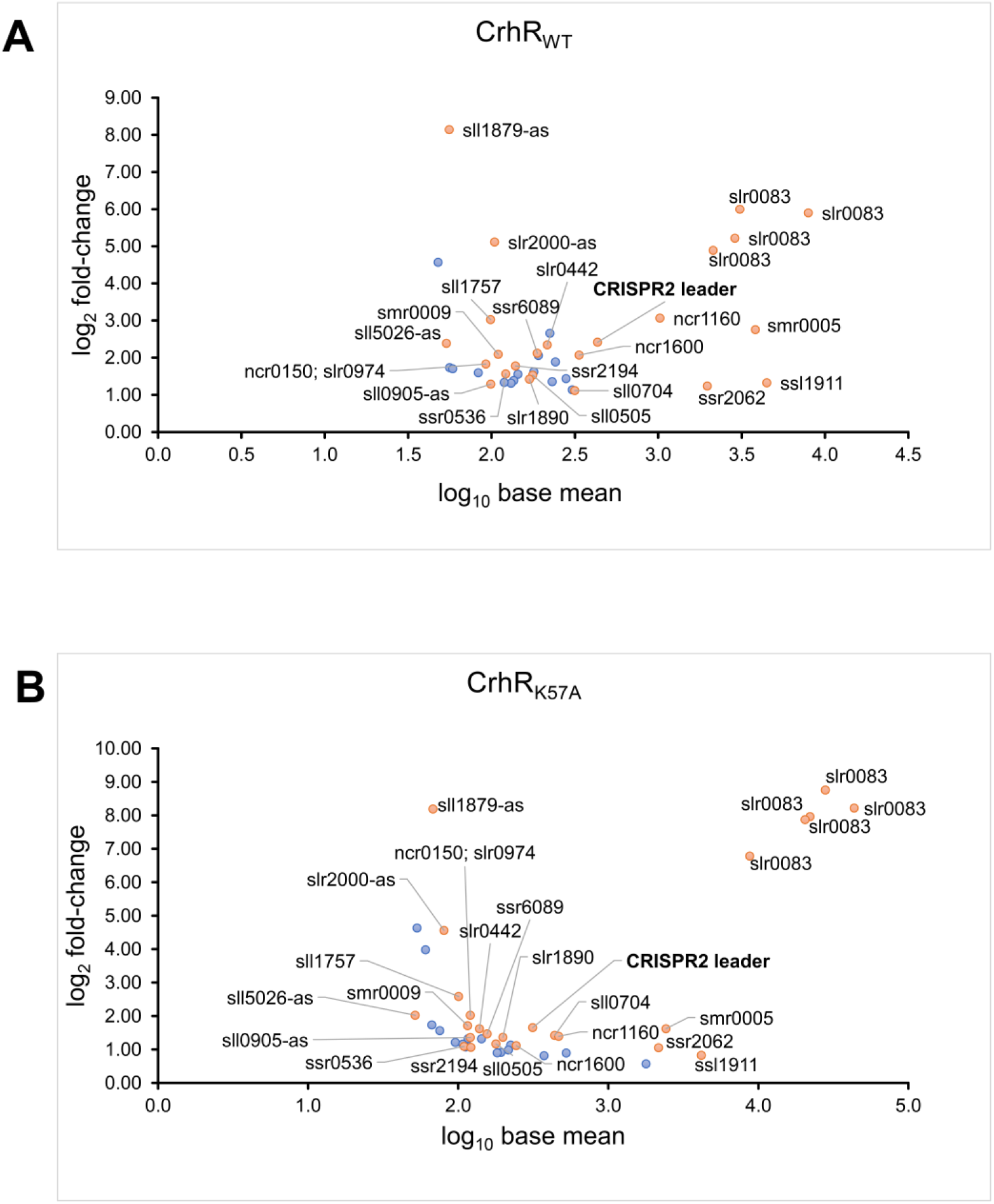
RNA target enrichment in the *in vitro* RNA pulldown from *Synechocystis* 6803. **A.** Recombinant His-tagged CrhR was used. **B.** The CrhR_K57A_ RNA helicase mutant was used. The peaks identified with PEAKachu are shown in the MA plot. Based on two biological replicates, 39 CrhR and 41 CrhR_K57A_ peaks were identified as significantly enriched (padj < 0.05, log_2_FC > 0. Common peaks for CrhR and CrhR_K57A_ are ivory-colored, and the unique peaks are shown in blue.

**Table 1.** RNA enriched from *Synechocystis* 6803 by *in vitro* pulldown using recombinant CrhR as bait. The experiment was performed in biological duplicates. Acronyms: asRNA, antisense RNA; Chr, chromosome.

| Peak start | Peak end | S | log<sub>2</sub>FC | p<sub>adj</sub> | Annotation | Description |
| --- | --- | --- | --- | --- | --- | --- |
| **Plasmid pSYSA** |  |  |  |  |  |  |
| 69586 | 69663 | + | 4.57 | 1.25E-07 | CRISPR2 array | S15R16 |
| 68651 | 68827 | + | 2.66 | 9.90E-08 | CRISPR2 array | R3S3R4S4R5 |
| 68370 | 68531 | + | 2.42 | 3.47E-09 | CRISPR2 leader | Transcribed leader of subtype III-Dv CRISPR Cas system and R1 |
| 69040 | 69125 | + | 1.89 | 1.56E-04 | CRISPR2 array | R7S7R8 |
| 68828 | 68996 | + | 1.44 | 3.08E-03 | CRISPR2 array | S5R6S6R7 |
| 69207 | 69355 | + | 1.33 | 2.71E-02 | CRISPR2 array | S10R11S11R12 |
| **Plasmid pSYSM** |  |  |  |  |  |  |
| 28098 | 28318 | + | 2.38 | 1.42E-03 | slI5026-as | asRNA to DNA phosphorothioation-associated putative methyltransferase |
| **Plasmid pSYSX** |  |  |  |  |  |  |
| 82429 | 82556 | + | 2.12 | 6.29E-05 | ssr6089-gene | Small 72 aa hypothetical protein |
| 27704 | 27866 | + | 1.36 | 1.58E-02 | ssr6030-gene | ORF and 3' UTR of small 72 aa hypothetical protein |
| **Chr** |  |  |  |  |  |  |
| 2888522 | 2888828 | + | 5.99 | 9.24E-54 | slr0083-gene | DEAD-box RNA helicase CrhR |
| 2887947 | 2888283 | + | 5.90 | 2.20E-60 | slr0083-gene | DEAD-box RNA helicase CrhR |
| 2888829 | 2889112 | + | 5.22 | 3.52E-45 | slr0083-gene | DEAD-box RNA helicase CrhR |
| 1446281 | 1446359 | - | 5.11 | 1.26E-10 | slr2000-as | asRNA to the gene of hypothetical protein |
| 2887606 | 2887893 | + | 4.89 | 2.76E-42 | slr0083-gene | DEAD-box RNA helicase CrhR |
| 2347288 | 2347369 | + | 3.06 | 5.96E-17 | ncr1160-ncRNA | Ncr1160 (SyR11) |
| 1242318 | 1242389 | - | 3.02 | 4.28E-07 | slI1757-5'UTR | asRNA to gene of hypothetical protein |
| 467144 | 467292 | + | 2.75 | 8.07E-15 | smr0005-5'UTR, gene | 5'UTR and ORF of photosystem I subunit XII |
| 2082308 | 2082609 | + | 2.34 | 6.55E-07 | slr0442-gene | Hypothetical protein |
| 1167431 | 1167719 | + | 2.09 | 2.10E-04 | smr0009-gene | Photosystem II PsbN protein |
| 3250506 | 3250676 | + | 2.07 | 2.25E-06 | ncr1600-ncRNA | 5'UTR of *rnj* encoding RNase J |
| 991513 | 991704 | - | 2.06 | 3.55E-05 | slI1463-gene | ATP-dependent zinc metalloprotease FtsH4 |
| 342867 | 343104 | + | 1.83 | 3.15E-03 | ncr0150-ncRNA; slr0974-gene | Nr0150; ORF of initiation factor IF-3 (*infC*) |
| 1654914 | 1655020 | + | 1.78 | 9.43E-04 | ssr2194-gene | Hypothetical protein |
| 1997946 | 1998013 | - | 1.73 | 2.26E-02 | slr0935-as-<br>asRNA | asRNA to gene encoding hypothetical<br>protein |
| 2839588 | 2839742 | + | 1.70 | 2.20E-02 | ssr1407-gene | Gene encoding hypothetical protein |
| 1141835 | 1141991 | - | 1.62 | 1.69E-03 | ssl1633-5'UTR, gene | 5'UTR and ORF of high light-inducible<br>polypeptide HliC |
| 1558635 | 1558806 | - | 1.60 | 1.54E-02 | ssl3769-gene;<br>ncl0710-ncRNA | Hypothetical protein; Ncl0710 |
| 2290635 | 2290796 | + | 1.56 | 6.75E-03 | ssr0536-5'UTR, gene | 5'UTR and ORF of hypothetical<br>protein |
| 3516172 | 3516339 | - | 1.56 | 4.78E-03 | slI0430-gene | HtpG, heat shock protein 90,<br>molecular chaperone |
| 3204827 | 3204897 | - | 1.53 | 6.96E-03 | slI0505-gene | Hypothetical protein |
| 822850 | 822979 | + | 1.42 | 8.23E-03 | slr1890-gene | ORF of bacterioferritin |
| 2082626 | 2082803 | + | 1.39 | 1.59E-02 | slr0442-gene | Hypothetical protein |
| 631992 | 632169 | - | 1.33 | 9.24E-04 | ssl1911-5'UTR, gene | 5'UTR and ORF of glutamine<br>synthetase inactivating factor IF7 |
| 1905897 | 1906083 | - | 1.31 | 3.45E-02 | slI1434-5'UTR, gene | 5'UTR and ORF of hypothetical<br>protein |
| 2028142 | 2028326 | + | 1.29 | 4.39E-02 | slI0905-as-<br>asRNA | asRNA to gene *maf* |
| 1052427 | 1052589 | + | 1.24 | 3.21E-03 | ssr2062-5'UTR, gene | 5'UTR and ORF of hypothetical<br>protein |
| 2270222 | 2270394 | - | 1.14 | 2.26E-02 | ncl1130-ncRNA | Ncl1130 |
| 117577 | 117740 | - | 1.12 | 2.65E-02 | slI0704-5'UTR, gene | 5'UTR and ORF of cysteine<br>desulfurase |

**Table 2.** RNA enriched from *Synechocystis* 6803 in an *in vitro* pulldown assay using recombinant His-CrhR_K57A_ as bait. The experiment was performed in biological duplicates.

| Replicon | Peak start | Peak end | Strand | log2 FC | padj | Annotation | Description |
| --- | --- | --- | --- | --- | --- | --- | --- |
| **Plasmid pSYSA** |  |  |  |  |  |  |  |
| pSYSA | 68370 | 68525 | + | 1.66 | 5.47E-09 | CRISPR2 leader | Transcribed leader of the subtype III-Dv CRISPR Cas system |
| pSYSA | 100746 | 101032 | + | 1.07 | 0.045406 | slr7104-gene | 5'UTR and ORF of transposase |
| **Plasmid pSYSM** |  |  |  |  |  |  |  |
| pSYSM | 28097 | 28309 | + | 2.03 | 0.005485 | sll5026-as | asRNA to DNA phosphorothioation-associated putative methyltransferase |
| pSYSM | 81234 | 81442 | + | 1.33 | 0.006107 | slr5082-gene | ORF of Rpn family recombination-promoting nuclease/putative transposase |
| pSYSM | 108252 | 108558 | + | 1.21 | 0.032833 | slr5118-gene | ORF of Rpn family recombination-promoting nuclease/putative transposase |
| pSYSM | 112774 | 113051 | + | 1.16 | 0.033782 | slr5124-gene | ORF of Rpn family recombination-promoting nuclease/putative transposase |
| **Plasmid pSYSX** |  |  |  |  |  |  |  |
| pSYSX | 82424 | 82546 | + | 1.47 | 0.000739 | ssr6089 | Small 72 aa hypothetical protein |
| **Chr** |  |  |  |  |  |  |  |
| NC_000911 | 2888558 | 2888865 | + | 8.76 | 0 | slr0083-gene | DEAD-box RNA helicase CrhR |
| NC_000911 | 2888152 | 2888426 | + | 8.22 | 0 | slr0083-gene | DEAD-box RNA helicase CrhR |
| NC_000911 | 1796717 | 1796808 | + | 8.19 | 3.42E-06 | sll1879-as | asRNA to the gene of two-component response regulator ycf55 |
| NC_000911 | 2887878 | 2888146 | + | 7.96 | 0 | slr0083-gene | DEAD-box RNA helicase CrhR |
| NC_000911 | 2888867 | 2889139 | + | 7.87 | 0 | slr0083-gene, 3'UTR | ORF and 3'UTR of CrhR |
| NC_000911 | 2887594 | 2887822 | + | 6.78 | 0 | slr0083-gene | DEAD-box RNA helicase CrhR |
| NC_000911 | 2315608 | 2315718 | + | 4.63 | 5.47E-09 | sll0169-as | asRNA to the gene of cell division protein Ftn2 homolog |
| NC_000911 | 1446283 | 1446373 | - | 4.56 | 1.49E-12 | slr2000-as | asRNA to gene of hypothetical protein |
| NC_000911 | 45835 | 46078 | - | 3.98 | 5.47E-09 | slr1494-as | asRNA to the gene of putative multidrug resistance family ABC transporter |
| NC_000911 | 1242313 | 1242398 | - | 2.58 | 2.87E-08 | sll1757-5'UTR | ORF of hypothetical protein |
| NC_000911 | 342865 | 343096 | + | 2.02 | 2.34E-05 | ncr0150-ncRNA; slr0974-gene | Ncr0150; ORF of initiation factor IF-3 (InfC) |
| NC_000911 | 2732766 | 2732918 | - | 1.74 | 0.009472 | sll0184-5'UTR, gene | 5'UTR and ORF of group 2 RNA polymerase sigma factor SigC |
| NC_000911 | 1167395 | 1167724 | + | 1.71 | 0.000146 | smr0009-gene | ORF of photosystem II PsbN protein |
| NC_000911 | 467114 | 467324 | + | 1.62 | 6.96E-22 | smr0005-5'UTR, gene | 5'UTR and ORF of photosystem I subunit XII |
| NC_000911 | 2082349 | 2082581 | + | 1.62 | 0.000109 | slr0442-gene | ORF of hypothetical protein |
| NC_000911 | 1807144 | 1807311 | - | 1.56 | 0.010578 | sll1873-5'UTR, gene | 5'UTR and ORF of hypothetical protein |
| NC_000911 | 117580 | 117742 | - | 1.42 | 1.54E-08 | sll0704-5'UTR, gene | 5'UTR and ORF of cysteine desulfurase |
| NC_000911 | 2347290 | 2347369 | + | 1.39 | 2.70E-06 | ncr1160-ncRNA | Ncr1160 |
| NC_000911 | 822850 | 823003 | + | 1.37 | 0.000606 | slr1890-gene | ORF of bacterioferritin |
| NC_000911 | 2028141 | 2028320 | + | 1.36 | 0.005349 | sll0905-as | asRNA to the gene maf |
| NC_000911 | 336220 | 336557 | - | 1.32 | 0.002837 | sll0933-gene; ssl1784-gene | ORF of hypothetical protein; ORF of 30S ribosomal protein S15 |
| NC_000911 | 3204824 | 3204940 | - | 1.16 | 0.004903 | sll0505-gene | ORF of hypothetical protein |
| NC_000911 | 1919637 | 1919895 | - | 1.13 | 0.004242 | sll1096-5'UTR, gene | 5'UTR and ORF of 30S ribosomal protein S12 |
| NC_000911 | 3250459 | 3250679 | + | 1.11 | 0.001265 | ncr1600-ncRNA | Ncr1600 (5'UTR of rnj) |
| NC_000911 | 2290587 | 2290787 | + | 1.09 | 0.041919 | ssr0536-5'UTR, gene | 5' UTR and ORF of hypothetical protein (84 aa) |
| NC_000911 | 1654915 | 1655063 | + | 1.06 | 0.039957 | ssr2194-gene | Small 65 aa hypothetical protein |
| NC_000911 | 1052426 | 1052590 | + | 1.05 | 5.10E-08 | ssr2062-5'UTR, gene | 5'UTR and ORF of hypothetical protein |
| NC_000911 | 1211732 | 1211980 | - | 0.99 | 0.009472 | sll1774-gene | ORF of 260 aa hypothetical protein |
| NC_000911 | 2224954 | 2225263 | - | 0.91 | 0.031283 | sll1863-gene | ORF of 107 aa hypothetical protein |
| NC_000911 | 69119 | 69384 | + | 0.90 | 0.036436 | slr1119-gene | ORF of 233 aa hypothetical protein |
| NC_000911 | 2644699 | 2644967 | + | 0.89 | 0.000729 | slr0628-5'UTR, gene | 5'UTR and ORF of *rpsN*, 30S ribosomal protein S14 |
| NC_000911 | 632000 | 632168 | - | 0.83 | 9.03E-05 | ssl1911-5'UTR, gene | 5'UTR and ORF of glutamine synthetase inactivating factor IF7 |
| NC_000911 | 862001 | 862259 | - | 0.81 | 0.011675 | ssl2233-5'UTR, gene | 5'UTR and ORF of *rpsT*, 30S ribosomal protein S20 |
| NC_000911 | 458684 | 458969 | - | 0.57 | 0.014432 | sll1515-gene | Orf of 149 aa hypothetical protein |

The most highly enriched transcripts for CrhR_K57A_ were asRNAs to the genes *sll0169*, *slr2000* and *slr1494*, which encode the DUF4101 and DnaJ-domain-containing protein Sll0169, the S-layer homology domain-containing protein Slr2000 and an ABC transporter subunit, respectively (**Table 2**).

Because the CRISPR2 leader RNA was enriched in co-IPs with both proteins, EMSA was performed to validate the interactions. For this purpose, the CRISPR2 leader RNA was synthesized by T7 RNA polymerase *in vitro* and used as an RNA substrate. For transcript synthesis, a DNA fragment with coordinates 68373-68498 on pSYSA was amplified using the primers EMSA_CRISPR2LeadeR-T7_Fw (which carries a T7 promoter sequence followed by two Gs) and EMSA_CRISPR2LeadeR-T7_Rv. The resulting 128 nt transcript was labeled with Cy3. Binding of 2 pmol of Cy3-labeled transcripts to various amounts of purified recombinant His-tagged CrhR or CrhR_K57A_, ranging from 1 to 50 pmol, was performed in the presence of poly(dI-dC) in high molar excess to the transcripts as a competitor to confirm the specificity of the RNA‒protein interaction. A gel shift of the CRISPR2 leader was observed upon the addition of only 1 pmol of CrhR (**Figure 5**). We concluded that the CRISPR2 leader transcript was strongly bound by both CrhR and CrhR_K57A_.

**Figure 5.**
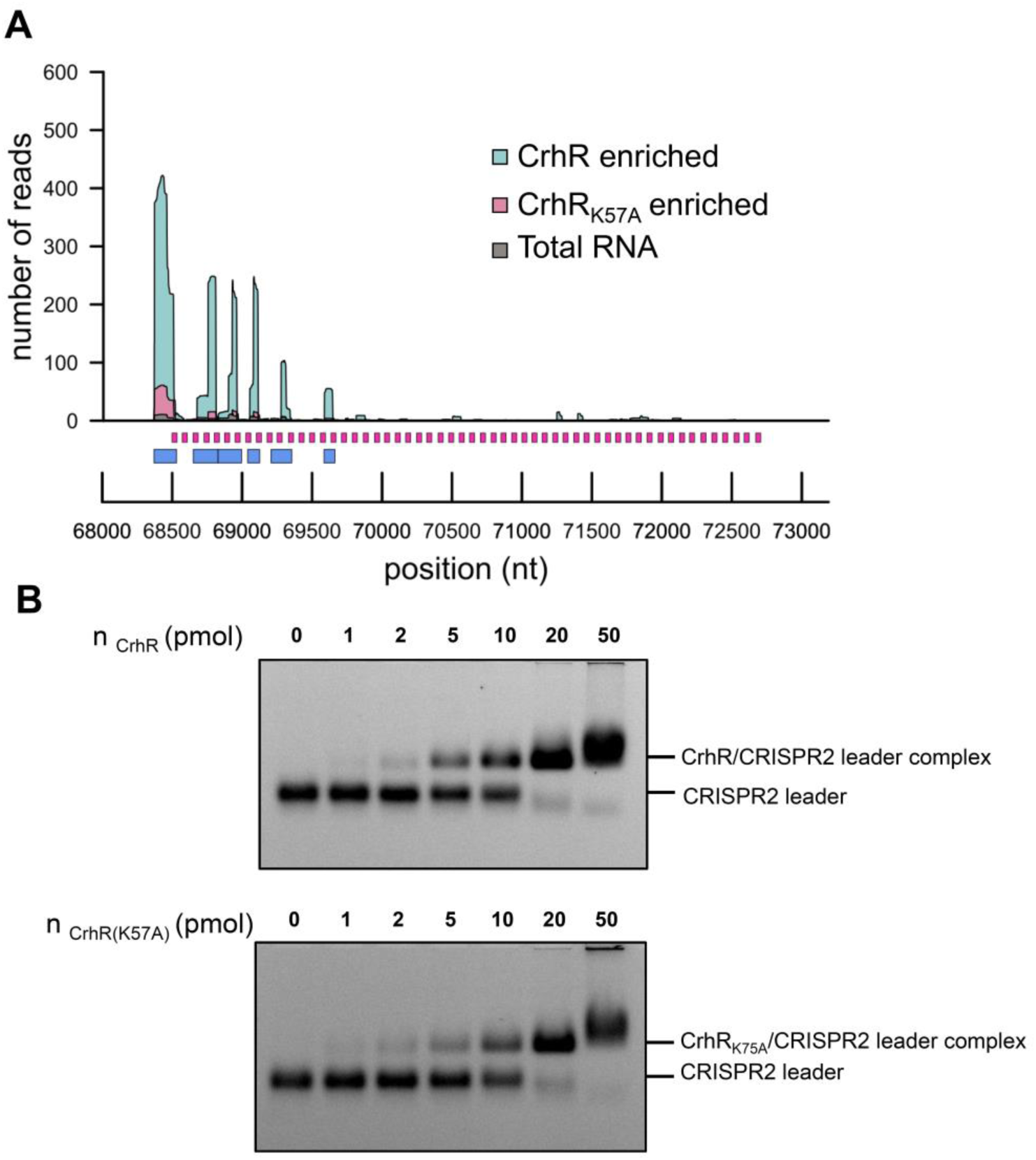
Interaction of the CrhR wild type and K57A mutant with the CRISPR2 leader. **A.** Enrichment of the CRISPR2 leader in the *in vitro* RNA pulldown from *Synechocystis* 6803 using recombinant His-tagged CrhR and CrhR_K57A_. Colored plots show the read coverage of the enriched RNAs. The experiment was performed in two biological replicates. CRISPR2 repeats and identified peaks are represented by magenta and blue boxes, respectively. **B.** EMSA showing the binding of CrhR (upper panel) and CrhR_K57A_ (lower panel) to the CRISPR2 leader RNA. EMSAs were performed with 2 pmol (81 ng) Cy3-labeled CRISPR2 leader RNA and the indicated amounts of purified His-tagged CrhR or CrhR_K57A_ in the presence of 1 μg of the competitor poly(dI-dC). Representative results from two independent experiments are shown.

### Effect of the Δ*crhR* mutation and redox stress conditions on CRISPR2 leader and crRNA accumulation

We next studied the effect of environmental stress conditions on *Synechocystis* 6803 wild type and the Δ*crhR* mutant. The cells were cultivated under standard growth conditions (low light and 30 °C) and exposed to either 20 °C or high light, followed by recovery under low light. Total RNA was extracted and hybridized with probes against the CRISPR2 leader or spacers 1-4. When testing the wild type and the mutant under standard and cold conditions (**Figure 6A**), we observed a lower level of CRISPR2 leader accumulation in Δ*crhR* than in the wild type. We analyzed the signal intensities normalized to those of 5S rRNA and observed that in the wild-type strain, the leader transcript intensity decreased by approximately 40% at 20 °C (**Figure 6B**). With respect to Δ*crhR*, we observed similar amounts of CRISPR2 leader transcripts under both conditions but generally lower amounts than in the wild-type. The accumulation of the CRISPR2 leader transcript did not seem to be affected by the change in temperature in the Δ*crhR* strain. These results indicated that CrhR, on the one hand, had a basal stabilizing effect on CRISPR2 leader transcript accumulation but that it had a destabilizing effect during temperature downshifts.

**Figure 6.**
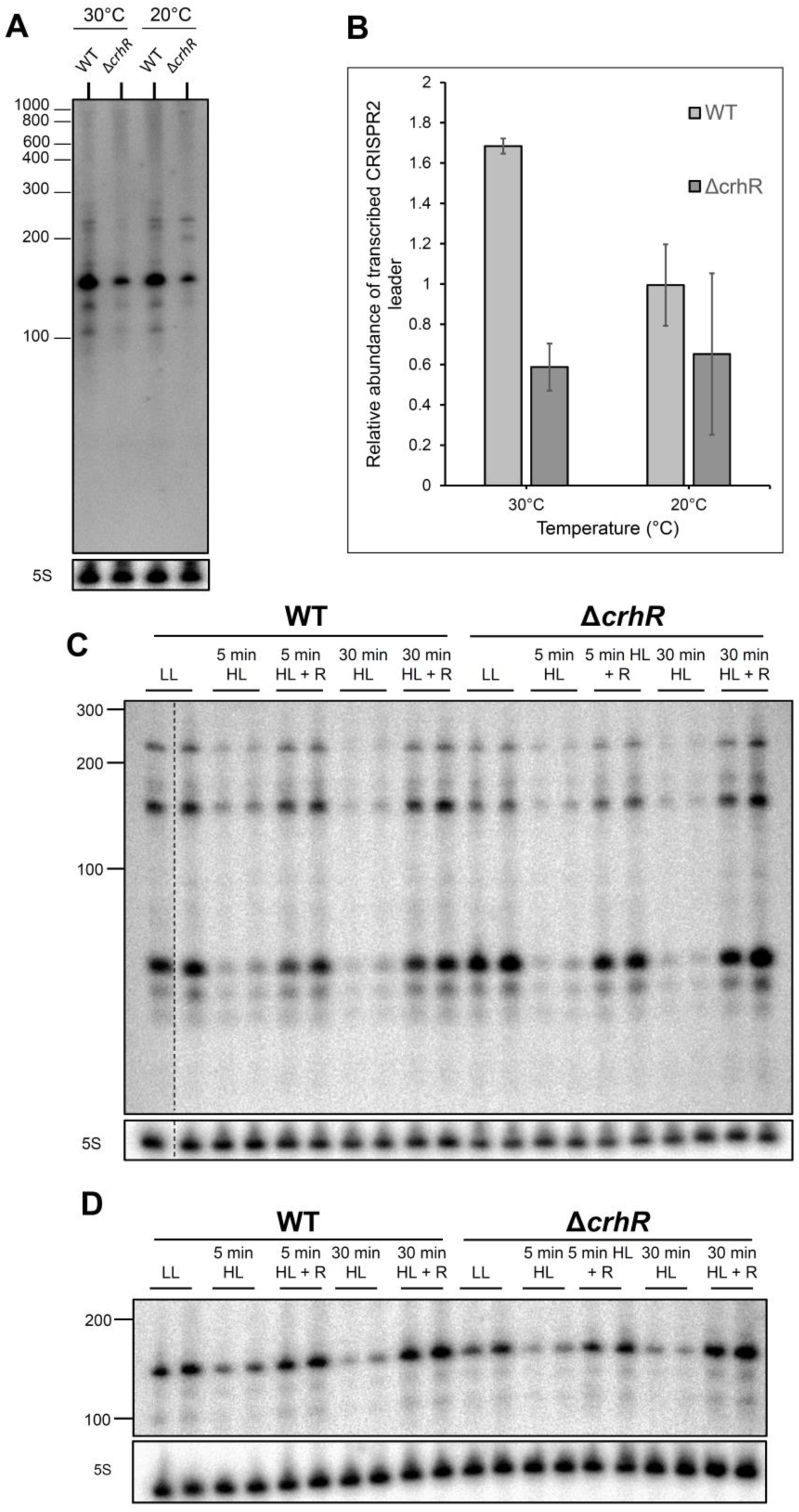
CRISPR2 leader and repeat-spacer accumulation in *Synechocystis* 6803 wild type and Δ*crhR* under cold and light stress conditions. **A.** Impact of *crhR* deletion on CRISPR2 leader transcript accumulation. The wild-type and Δ*crhR* were cultivated at 30 °C or incubated at 20 °C for 2 h. Total RNA was extracted, and 10 µg was loaded per lane on an 8 M urea-10% PAA gel. The strains were tested for accumulation of the CRISPR2 leader transcript by hybridization using a transcript oligonucleotide. Hybridization of 5S rRNA is shown as the control for equal loading. A representative of biological duplicates is shown. **B.** Relative amounts of CRISPR2 leader transcripts were normalized to 5S rRNA intensity and quantified. Ten µg of *Synechocystis* wild type and Δ*crhR* cultivated under low light (LL) or high light (HL) conditions, with or without recovery (R), were separated on a 10% denaturing polyacrylamide gel. A ^32^P-labeled transcript probe specific to **C**. spacer1– spacer4 or **D**. CRISPR2 leader was hybridized. Hybridization of 5S rRNA is shown for the control of equal loading.

When testing the influence of high light on CRISPR2 leader and repeat spacer array transcript accumulation, we observed a rapid decrease in the accumulation of both transcripts after exposure to high light for 5 min (**Figure 6C** and **D**). The same observations were made after 30 min under high-light conditions. For recovery, the cultures were again exposed to low light, and cultivation was continued for 2 h. After the recovery phase, the number of leader and repeat-spacer transcripts was similar to that before high light exposure. These results suggest rapid degradation of the leader and spacer transcripts by an unknown mechanism and rapid adaptation to changes in the redox status of the cell.

### Determination of CrhR amino acid residues interacting with the CRISPR2 leader

To confirm the interaction unambiguously and to identify the amino acid residues of CrhR that interact with the CRISPR2 leader, CrhR was cross-linked to the CRISPR2 leader RNA *in vitro*. In total, we obtained 12 cross-linked peptide fragments for two replicates using UV and CrhR_K57A_, one for CrhR and 3 for CrhR_K57A_ using the chemical cross-linker 1,2,3,4-diepoxybutane (**Figure 7A**). The amino acid residues cross-linked to the RNA were determined as described previously^46^ and are shown in **Figure 7B**. None of the cross-linked amino acid residues were located within the known conserved motifs of DEAD-box RNA helicases (**Figure S4**). We used AlphaFold 2^47,48^ to predict the three-dimensional structure of CrhR. Consistent with recent reports that CrhR exists in solution predominantly as a homodimer^49^, AlphaFold modeled it as a dimer and predicted alpha helices and beta folds in the most conserved part of the protein. No structure was predicted for the C-terminal section of the protein, consistent with a lack of sequence conservation (**Figure 7C**). The dimeric structure of these proteins is consistent with the homodimeric structure of the *Geobacillus stearothermophilus* RNA helicase CshA, the closest homolog of CrhR (43.57% sequence identity), for which the structure has been resolved^50^. By analyzing the model, we found that the amino acid residues L103, F104, H225, and C371 were located on the surface of CrhR, whereas the amino acid residue C184 was not. We could not draw a conclusion about the possible location of the amino acid residue P443 because the modeling failed for the 65 C-terminal residues.

**Figure 7.**
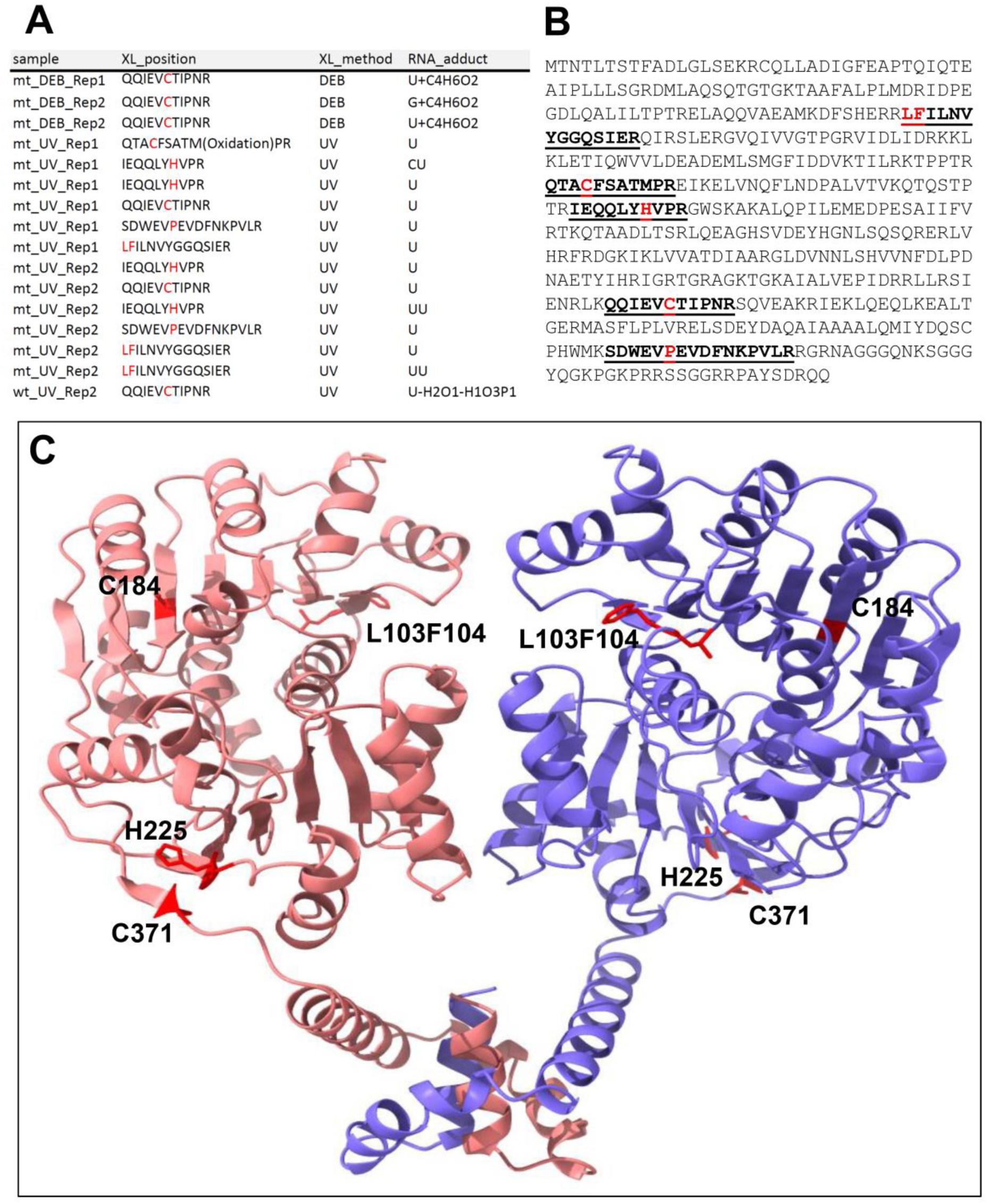
Crosslinking of CrhR to CRISPR2 leader RNA. **A.** Overview of the peptide fragments and RNA adducts detected in two replicate samples harboring CrhR_K57A_ (mt) or CrhR (wt). **B.** Sequence of CrhR. The amino acid residues of CrhR_K57A_ cross-linked by UV treatment at 254 nm to the CRISPR2 leader are highlighted in red. The respective detected peptide fragments are underlined and in boldface letters. The QQIEVcTIPNR peptide was also detected for CrhR and in addition by chemical cross-linking using 1,2,3,4-diepoxybutane^77^ instead of UV treatment. **C.** Structure of a CrhR homodimer (amino acids 9 to 427) predicted by Alphafold2^47,48^. The CrhR amino acid residues cross-linked to the CRISPR2 leader transcript and identified by LC‒MS are highlighted in red. The cross-linked CrhR residues in the context of conserved sequence segments and previously identified functionally relevant domains are given in **Figure S4**.

## Discussion

The main role of CRISPR/Cas systems is defense against encountered phages. Therefore, their constitutive expression might be expected. However, there is mounting evidence that some CRISPR/Cas systems, such as those in *E. coli*^7–10^ and *Sulfolobus islandicus*^11,12^, are regulated at the transcriptional level. When the risk of phage infection is high, *Pseudomonas aeruginosa* is regulated by environmental factors, such as temperature or high cell density; LasI/R and RhlI/R, two autoinducer pairs from the quorum sensing pathway, promote the expression of the type I-F CRISPR/Cas system^15,16^. Furthermore, resource availability can strongly influence *cas* gene expression. The cAMP receptor protein (CRP) binds to DNA in the presence of its co-factor cAMP, the level of which depends on the availability of glucose in the environment. In the phytopathogen *Pectobacterium atrosepticum*, CRP increases the expression of type I-F *cas* genes when glucose is scarce^14^, whereas *cas* transcription is negatively regulated by the cAMP-CRP complex in the type I-E system of *E. coli* when glucose is available^51^.

Here, we showed that RpaB, a DNA-binding response regulator, controls the transcription of the type III-Dv *cas* operon in the cyanobacterium *Synechocystis* 6803. RpaB is a redox-responsive transcription factor that is highly conserved in cyanobacteria and is a key regulator of light acclimation^52^. RpaB controls a large panel of genes relevant for photosynthesis, photoprotection, membrane transport^30^. Analysis of the distribution of the HLR1 binding motif of RpaB in *Synechocystis* 6803 showed that RpaB functions as an activator under low-light conditions when the HLR1 motif is located at positions -66 to -45 to the TSS and as a repressor if located elsewhere in the promoter^30^. The finding that the abundance of crRNAs for the III-Dv system in *Synechocystis* 6803 varies greatly between different environmental conditions can therefore be partially explained by the control of the *cas* gene promoter through the binding of RpaB to HLR1 at an activating position. The availability of Cas proteins can certainly limit the formation of Cas complexes and the protection of the crRNAs bound to them. However, we were puzzled that the repeat-spacer array promoter, albeit very strong, not only lacked an HLR1 motif but also exhibited slightly greater activity in reporter gene assays under high light, contrary to the *cas* gene promoter.

This led us to consider the interaction between CrhR and the 125 nt leader transcript^19^. The CRISPR leader is usually understood as a longer region containing the promoter^2,53^, regulatory sequence elements important for adaptation^54–56^ and the TSS of the repeat-spacer array. CRISPR leaders have mostly been studied for their roles in spacer acquisition in the genome. However, they may also play an important role in the posttranscriptional regulation of precrRNAs and affect crRNA maturation and interference. The sRNA-dependent posttranscriptional regulation of a CRISPR array was identified in *P. aeruginosa*, where binding of the sRNA PhrS to the leader of a type I-F system repressed the Rho-dependent termination of CRISPR array transcription^57^.

We showed that the CRISPR2 leader transcript also exists as a distinct sRNA in the cell and that the accumulation of the CRISPR2 leader and crRNAs is strongly affected by the cellular redox status. We found that this leader RNA was highly enriched in *in vitro* co-IPs with recombinant CrhR and CrhR_K57A_. We confirmed the leader-CrhR (and leader-CrhR_K57A_) interaction by EMSA and identified the interacting amino acid residues by protein‒RNA cross-linking coupled to mass spectrometry analysis^46,58^. The cross-linked residues L103/F104, H225, C371, C184 and P443 do not match positions previously described to be involved in the interactions between DEAD-box RNA helicases and their substrates^59^. However, these residues are in line with calculations of UV cross-linking efficiencies for different amino acids, which, among others, included phenylalanine (F), histidine (H) and proline (P), which were found here^60^. Moreover, the systematic analysis of interactions between mutagenized RNA and protein variants suggested that π-stacking interactions between aromatic amino acids (such as Y, F or H) and guanosine or uridine residues are important for cross-linking and for flanking amino acids^61^, whereas cysteine is prone to cross-linking due to its high reactivity^58^. Four of the 6 amino acids identified here matched these criteria, and L103 was flanked by aromatic amino acids on both sides (**Figure S4**). Moreover, with the exception of C184, these amino acids were all predicted to be on the surface of a dimeric CrhR model (**Figure 7C**). Thus, both aspects are consistent with the possible involvement of these residues in RNA recognition and binding and indicate the potential for further analyses in the future.

RNA helicases are enzymes that can modify RNA structures. Therefore, they are associated with all aspects of RNA metabolism, such as the regulation of gene expression, RNA maturation and decay, transcription and the packaging of RNA into ribonucleoprotein particles^62,63^, processes that are also relevant for the formation of CRISPR–Cas complexes. The expression of *crhR* is regulated by the redox status of the electron transport chain^31^ and becomes strongly enhanced in response to a decrease in temperature^64^. CrhR plays a role in the modulation of multiple metabolic pathways during cold acclimation^35^ and is indispensable for energy redistribution and the regulation of photosystem stoichiometry at low temperatures^65^. Consistent with these physiological functions, CrhR is localized to the thylakoid membrane but also cosedimented with degradosome and polysome complexes^66^. Our data showed decreased leader and crRNA accumulation upon shifts to high light or low nitrogen, which was most pronounced upon addition of the inhibitor DBMIB, suggesting that these conditions constitute redox stress effects. A redox component involved in the expression of CRISPR‒Cas systems has not been previously shown. However, such regulation is highly important for cyanobacteria, which are the only prokaryotes that perform oxygenic photosynthesis. In fact, phage adsorption to the cyanobacterial host, replication, modulation of host cell metabolism, and survival in the environment following lysis all exhibited light-dependent components^67^.

Indeed, the transcriptional control of *cas* gene transcription through RpaB and the recruitment of the DEAD-box RNA helicase CrhR by the leader transcript are consistent with this mode of regulation (**Figure 8**). The involvement of CrhR in this process adds to recent reports on the connection between components of the degradosome and the type III CRISPR‒Cas machinery^22,68^. Our results are furthermore consistent with recent results of unbiased screens that multiple host genes can affect CRISPR expression^13^. The here described parallel control of *cas* gene transcription by the transcription factor RpaB and the effect of CrhR on CRISPR leader and crRNA accumulation highlight the intriguing complexity of CRISPR‒Cas regulation in the context of the host cell.

**Figure 8.**
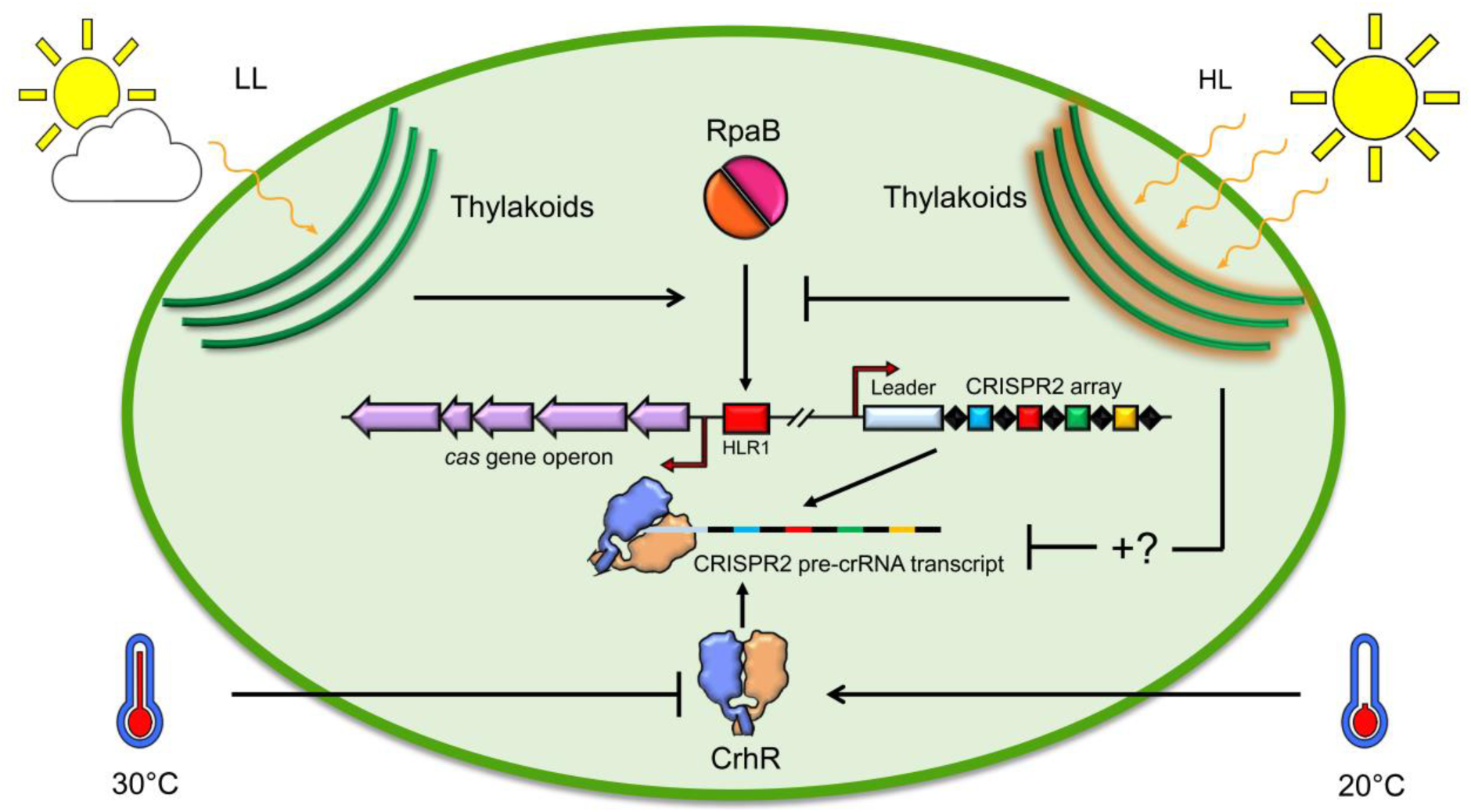
Multilevel redox control of CRISPR2 expression. The transcription factor RpaB binds to its HLR1 motif (red rectangle) under low light, initiating the expression of the *cas* gene operon (purple arrow), which encodes the effector complex of the type III-Dv system in *Synechocystis* 6803. Under high light conditions, the change in the redox status of the photosystems, located in the thylakoid membrane (green arcs), leads RpaB to dissociate from its HLR1 motif, resulting in repression of the transcription of the *cas* gene operon (purple arrows). At the posttranscriptional level, high light conditions lead to a decrease in the CRISPR2 leader and repeat-spacer transcript accumulation by an unknown mechanism (question mark). The DEAD-box RNA helicase CrhR recognizes the leader transcript. The attachment of helicase to the CRISPR2 leader transcript is temperature-dependent. At low temperature (20 °C), CrhR binds to the leader transcript, whereas at higher temperature (30 °C), it inhibits this interaction. HL: high light; LL: low light; TM: thylakoid membrane.

## Materials and Methods

### Strains and growth conditions

Cultures of the wild type and different mutant strains of *Synechocystis* 6803^33,36^ were grown at 30 °C in liquid BG11 medium^69^ supplemented with 20 mM TES (N-[Tris-(hydroxymethyl)-methyl]-2-aminoethane sulfonic acid) under continuous illumination with white light at 50 μmol photons m^-2^ s^-1^ without shaking unless otherwise mentioned. The flasks were aerated with ambient air through a glass tube and a sterile filter for constant and fast growth. To induce gene expression from the Cu^2+^-responsive promoter P*_petE_*_70_, 2 μM CuSO_4_ was added to exponentially growing cells in BG11 medium without Cu_2+_.

Mutant strains of *Synechocystis* 6803 were grown in the presence of the appropriate antibiotics at the following concentrations: spectinomycin (sp) (20 μg/mL) and kanamycin (km) (50 μg/mL) for the Δ*crhR*/FLAG, Δ*crhR*/FLAG-CrhR, and Δ*crhR*/FLAG-CrhR_K57A_ strains. For the cold and high light stress experiments, *Synechocystis* 6803 strains were cultivated at 30 °C under continuous white light (30-50 µmol m^-2^ s^-1^) and shaken until OD_750nm_ = 0.6 was reached. For cold stress, the cultures were then split into two groups: one group was cultivated at 30 °C, and the second group was placed in a water bath and kept at 20 °C with ice for 2 h. Under high light conditions, if not otherwise indicated, the cells were exposed to 300 µmol m^-2^ s^-1^ for 5 or 30 min. For recovery, the cells were returned to low light conditions for 2 h. To construct *E. coli* strains for the expression of recombinant CrhR, the *crhR*_WT_ and *crhR*_K57A_ reading frames were cloned from the respective *Synechocystis* 6803 strains^33,36^ and inserted into the pQE-70 vector upstream of a segment encoding a 6×His-tag and subsequently transformed into *E. coli* M15. *E. coli* strains were grown in liquid LB media (10 g l^-1^ bacto-tryptone, 5 g l^-1^ bacto-yeast extract, 10 g l^-1^ NaCl) with continuous agitation or on agar-solidified (1.5% [w/v] Bacto agar) LB supplemented with appropriate antibiotics at 37 °C.

### RNA isolation

*Synechocystis* 6803 cells were collected by vacuum filtration through hydrophilic polyethersulfone filters (Pall Supor®-800, 0.8 μm), transferred to a tube containing 1 mL of PGTX buffer^71^, snap-frozen in liquid nitrogen and stored at -80 °C until further use. RNA was extracted as described previously^36^, and the RNA concentration was determined using a NanoDrop ND-1000 spectrophotometer (Peqlab).

### Recombinant protein expression and purification

*E. coli* M15 was transformed with the vectors pQE70:crhR-6×His and pQE70:crhR_K57A_-6×His for overexpression of the recombinant His-tagged proteins CrhR and CrhR_K57A_. Overnight cultures were diluted 1:100 in fresh LB medium supplemented with ampicillin and kanamycin and grown to an OD_600_ of 0.7. Protein expression was induced by adding isopropyl-β-D-thiogalactopyranoside (IPTG; 1 mM final concentration). Three hours after IPTG induction, the cells were harvested by centrifugation at 6,000 × g for 10 min at room temperature. The cell pellets were resuspended in lysis buffer (50 mM NaH_2_PO_4_ (pH 8), 1 M NaCl, 10% glycerol, 15 mM imidazole, and cOmplete™ Protease Inhibitor Cocktail (Roche)) and lysed using the One Shot Constant Cell Disruption System (Constant Systems Limited, United Kingdom) at 2.4 kbar. Cell debris was pelleted by centrifugation at 13,000 × g for 30 min at 4 °C, and the lysate was filtered through 0.45 µm Supor-450 filters (Pall). Recombinant proteins were immobilized on a HiTrap Talon crude 1 mL column (GE Healthcare), equilibrated with buffer A (50 mM NaH_2_PO_4_ (pH 8), 500 mM NaCl), and eluted with elution buffer B (50 mM NaH_2_PO_4_ (pH 8), 500 mM imidazole, 500 mM NaCl).

Recombinant 6×His-tagged RpaB in Rosetta (DE3)+pLysS was expressed as described^39^, using phosphate-buffered TB medium instead of 2 × YT medium. RpaB was purified following the same protocol^39^ using Precellys 24 (Bertin Technologies) to disrupt the cells and 0.5 mL of bedded Ni-NTA agarose beads (Qiagen GmbH) to bind 6×His-tagged proteins. The protein concentration was calculated using the Bradford assay. The protein samples were mixed with Coomassie Plus (Bradford) Assay Reagent (Thermo Fisher Scientific) in a 96-well plate. The absorption at 595 nm was measured using a Victor^3^ 1420 multilabel plate reader (Perkin Elmer). The protein concentration was calculated based on a bovine serum albumin calibration curve.

### In vitro His-tag affinity purification and RNA pulldown

Recombinant proteins were isolated from *E. coli* M15 strains via precipitation on Dynabeads™ magnetic beads (125 μl), which bind histidine-tagged proteins. To prove the coupling of His-tagged proteins to the beads, an aliquot (5%) of a sample containing beads was washed after protein pulldown from *E. coli* M15 and used for SDS‒PAGE analysis. The beads coupled with the His-tagged protein were further incubated in 25 mM Tris-HCl RNA elution buffer containing 2 M NaCl to eliminate contaminating RNA molecules from *E. coli*, washed in 1× TBS, and incubated in the cell lysate of *Synechocystis* 6803 wild type for 20 min. RNA from wild-type *Synechocystis* coprecipitated with the recombinant His-tagged proteins was eluted in the same RNA elution buffer and further utilized for the generation of libraries for Illumina sequencing.

### CrhR-CRISPR2 leader RNA cross-linking and enrichment of cross-linked peptide-RNA heteroconjugates

We used 10 min of UV irradiation at 254 nm to covalently cross-link approximately 1 nmol of the complex formed between CRISPR2 leader RNA and the CrhR protein in a volume of 100 µL in buffer containing 50 mM NaH_2_PO_4_, 300 mM NaCl, and 250 mM imidazole (pH 8.0) as described previously^22^. Subsequently, cross-linked peptide‒ RNA heteroconjugates were enriched according to our previously established workflow^46,58^. We ethanol-precipitated the samples and resuspended the pellet in buffer containing 4 M urea and 50 mM Tris-HCl (pH 7.9). The urea concentration was subsequently decreased to 1 M by adding 5 vol of 50 mM Tris-HCl (pH 7.9). The RNA was hydrolyzed by adding 1 µg of RNase A and T1 (Ambion, Applied Biosystems) at 52 °C for 2 h, followed by digestion with benzonase at 37 °C for 1 h and trypsin (Promega) digestion overnight at the same temperature. To remove the non-cross-linked RNA fragments and to desalt the sample, the sample was passed through a C18 column (Dr. Maisch GmbH), followed by enrichment of the cross-linked peptides over an in TiO_2_ column (GL Sciences) according to existing protocols^46^ but using 10 µm TiO_2_ beads as described previously^22^. The samples were subsequently dried, resuspended in 5% v/v acetonitrile and 1% v/v formic acid, and subjected to liquid chromatography and mass spectrometry analysis.

### Analysis by mass spectrometry

A nanoliquid chromatography system (Dionex, Ultimate 3000, Thermo Fisher Scientific) coupled with a Q Exactive HF instrument (Thermo Fisher Scientific)^46^ was used for liquid chromatography and mass spectrometry analysis. Online ESI-MS was performed in data-dependent mode using the TOP20 HCD method. All precursor and fragment ions were scanned in the Orbitrap, and the resulting spectra were measured with high accuracy (< 5 ppm) at both the MS and MS/MS levels. A dedicated database search tool was used for data analysis^58^.

### Promoter activity assay

The promoter region and 5’UTR of the CRISPR2 *cas* gene operon was PCR-amplified with the primer pairs prom_*cas10*_luxAB_fw and prom_*cas10*_luxAB_rev to amplify the wild-type promoter and the primer pairs prom_*cas10*_mut_luxAB_fw and prom_*cas10*_luxAB_rev to substitute the ACAA motif in the conserved HLR1 site with a GGGG motif. For the CRISPR2 array promoter, we cloned the 100 base pair region upstream of the transcription start site (68274-68373) with the primers Prom_CRISPR2_fw and Prom_CRISPR2_RBS_rev to fuse the promoter with an artificial ribosome binding site^72^. The pILA backbone, containing a promoterless *luxAB* gene^38^, was amplified in three parts with the primer pairs pILA_1_fw (or pILA_1_RBS_fw for the CRISPR2 array promoter)/pILA_1_rev, pILA_2_fw/pILA_2_rev, and pILA_3_fw/pILA_3_rev. Primers were designed to overlap adjacent fragments. PCR fragments were assembled using AQUA cloning^73^ and transformed into *E. coli* DH5alpha. The resulting strains were named pILA-P_CRISPR2_cas10nat_ and pILA-P_CRISPR2_cas10mut_, respectively. The resulting constructs were subsequently transformed into an engineered *Synechocystis* 6803 strain, which carries the *luxCDE* operon encoding the enzymes for the synthesis of decanal^74^. Segregation of the constructs was achieved by transferring single clones to new BG11-0.75% Kobe Agar plates containing increasing concentrations of kanamycin (10-50 µg/mL). Full segregation was verified by PCR using the primers pIGA-fw and pIGA_rev and sequencing. The clones with segregated pILA constructs were grown in BG11 supplemented with 50 µg/µL kanamycin, 10 µg/µL chloramphenicol and 10 mM glucose under continuous light (30-50 μmol photons m^-2^ s^-1^) and shaken until they reached the mid-logarithmic phase (OD_750 nm_ = 0.7 to 0.8). Cultures were diluted to OD_750 nm_ = 0.4 prior to exposure to high light conditions (300 μmol photons m^-2^ s^-1^) for four hours. Afterward, the cells were placed back in low light (40-50 µmol m^-2^ s^-1^). As shown in **Figure 2C**, DCMU was added to the cells during the high-light phase (at a final concentration of 50 µM). As shown in **Figure 3B**, the cells were kept under high light after the initial 4 h of exposure and were not switched back to low light again. Bioluminescence was measured *in vivo* by using a VICTOR^3^ multiplate reader (PerkinElmer) at total light counts per second. Cell suspensions (100 µL) were measured in a white 96-well plate (CulturePlate^TM^-96, PerkinElmer). Bioluminescence was measured before and after exposure to high light and every 30-60 min during recovery under low light. Next, we exposed the cells to both high and low light. On the basis of the results of preliminary tests, we noticed that the cellular production of decanal was not sufficient for monitoring bioluminescence *in vivo*. Therefore, we added 2 µL of decanal prior to the measurements. A strain carrying the promotorless *luxAB* gene served as a negative control. A strain carrying the P_Syr9_::*luxAB* construct was used as a control strain. Technical triplicates were measured. Statistical relevance was calculated using a 2-tailed t test in Excel (Microsoft).

### Electrophoretic mobility shift assay (EMSA)

For the binding of RpaB to wild-type and mutated HLR1 motifs, regions of interest were PCR-amplified from the pILA-PCRISPR2_*cas10* construct using the primers EMSA_P*cas10*_HLR1_rev and EMSA_P*cas10*_HLR1_rev, which were labeled with Cyanine 3 (Cy3) at the 5’ end. The HLR1 motif from the *psbA2* promoter was used as a positive control and amplified using the primer pair EMSA_P*psbA2*_fw and EMSA_P*psbA2*_rev. Different amounts of eluted 6×His-tagged RpaB (0-250 pmol) were mixed with 0.5 pmol of Cy3-labeled DNA target in binding buffer (20 mM HEPES-NaOH, pH 7.6; 40 mM KCl; 0.05 mg/mL BSA; 5% glycerol; 0.1 mM MnCl2; 1 mM DTT; 0.05 µg/µL poly(dIdC)). The reaction mixture was incubated for 30 min at room temperature in the dark. Electrophoresis was performed in a 3% agarose-0.5 × TBE gel. The gel was run for 60 min in the dark at 80 V and 4 °C. The signals were visualized with a Laser Scanner Typhoon FLA 9500 (GE Healthcare) using a green-light laser and Cy3 filter.

Binding of CrhR or CrhR_K57A_ to 0.2 pmol or 2 pmol of Cy3-labeled RNA was performed in buffer containing 20 mM HEPES-KOH (pH 8.3), 3 mM MgCl_2_, 1 mM DTT, and 500 μg/mL BSA. As a substrate competitor, 1 μg of LightShift poly(dIdC) (Thermo Fisher Scientific) was added. The reactions were incubated at room temperature for 15 min prior to loading on 2% agarose-TAE gels.

### Library preparation for RNA-seq

Total RNA was subjected to Turbo DNase (Thermo Fisher Scientific), purified, and size separated using an RNA Clean & Concentrator-5 Kit (Zymo Research) and treated with 5’-polyphosphatase (Epicenter) as described previously^36^. The RNA was phosphorylated at the 5’ end by T4 polynucleotide kinase (NEB) and ligated to the 5’ adapter (**Table S2**). A ThermoScript Reverse Transcriptase Kit (Invitrogen) was used for cDNA synthesis, and the cDNA was amplified with Phusion High-Fidelity DNA polymerase (Thermo Fisher Scientific) using PCR primers 1 and 2 (**Table S2**). The PCR conditions were 98 °C for 30 s, followed by 18 cycles of denaturation at 98 °C for 10 s, primer annealing at 60 °C for 30 s, and extension for 15 s at 72 °C, and a final extension step at 72 °C for 2 min. The ExoSAP-IT PCR Product Cleanup Reagent (Thermo Fisher Scientific) was used for primer removal, and the samples were further purified with the NucleoSpin® Gel and PCR Clean-up Kit and eluted with 20 µL of NE buffer. A 10 µL aliquot of each prepared DNA library was sequenced on an Illumina sequencer.

### RNA-seq data analysis

RNA-seq data analysis was performed using tools installed in usegalaxy.eu. The paired-end or single-end reads were trimmed, and adapters and reads shorter than 14 nt were filtered out by Cutadapt 1.16^75^. Mapping was performed on the chromosome and plasmids of *Synechocystis* 6803 by Bowtie2 2.3.4.3 with the parameters for paired-end reads: -I 0 -X 500 --fr --no-mixed --no-discordant --very-sensitive^43^. Unmapped reads were filtered. Peak calling of the mapped reads was performed using PEAKachu 0.1.0.2 with the parameters --pairwise_replicates --norm_method deseq --mad_multiplier 2.0 –fc_cutoff 1 --padj_threshold 0.05.

## Data availability

The RNA-seq data have been deposited in the SRA database https://www.ncbi.nlm.nih.gov/sra/ and are openly available under the accession numbers SRX6451369 to SRX6451374. All mass spectrometry proteomics datasets analyzed during this study are available in the Proteomics Identifications Database (PRIDE, at https://www.ebi.ac.uk/pride/) under the project accession number PXD047440.

## Acknowledgments

We thank Yukako Hihara, Saitama, Japan, and Jogadhenu S. S. Prakash, Hyderabad, India, for the *E. coli* RpaB overexpression and *Synechocystis ΔcrhR* strains, respectively. We thank Richard Reinhardt and his team, Cologne, Germany, for sequence analyses. The support of Sergey Moshkovskiy and Olexandr Dybkov in interpreting the mass spectrometry data is greatly appreciated. We thank Monika Raabe, Göttingen, Ingeborg Scholz and Viktoria Reimann, Freiburg, Germany, for their expert technical assistance and George Owttrim, Edmonton, Canada, for discussions about CrhR.

## AUTHOR CONTRIBUTIONS

W.R.H. designed the work. Protein‒RNA cross-linking experiments and identification of cross-linked peptide-RNA bonds were performed by A.W. and H.U. The analyses of RpaB effects, CRISPR2 leader and repeat-spacer accumulation were performed by R.B. All the other CrhR-related experiments were carried out by A.M. The construction of cDNA libraries and the analysis of the pull-down results was performed by A.M. and C.S. A.M., R.B. and W.R.H. wrote the paper with contributions from all the authors.

## References

1. Makarova, K.S., Grishin, N.V., Shabalina, S.A., Wolf, Y.I., and Koonin, E.V. (2006). A putative RNA-interference-based immune system in prokaryotes: computational analysis of the predicted enzymatic machinery, functional analogies with eukaryotic RNAi, and hypothetical mechanisms of action. Biol. Direct 1, 7.

2. Brouns, S.J.J., Jore, M.M., Lundgren, M., Westra, E.R., Slijkhuis, R.J.H., Snijders, A.P.L., Dickman, M.J., Makarova, K.S., Koonin, E.V., and Oost, J. van der (2008). Small CRISPR RNAs guide antiviral defense in prokaryotes. Science 321, 960– 964.

3. Makarova, K.S., Wolf, Y.I., Iranzo, J., Shmakov, S.A., Alkhnbashi, O.S., Brouns, S.J.J., Charpentier, E., Cheng, D., Haft, D.H., Horvath, P., et al. (2020). Evolutionary classification of CRISPR-Cas systems: a burst of class 2 and derived variants. Nat. Rev. Microbiol. 18, 67–83.

4. Makarova, K.S., Wolf, Y.I., Alkhnbashi, O.S., Costa, F., Shah, S.A., Saunders, S.J., Barrangou, R., Brouns, S.J.J., Charpentier, E., Haft, D.H., et al. (2015). An updated evolutionary classification of CRISPR-Cas systems. Nat. Rev. Microbiol. 13, 722–736.

5. Özcan, A., Krajeski, R., Ioannidi, E., Lee, B., Gardner, A., Makarova, K.S., Koonin, E.V., Abudayyeh, O.O., and Gootenberg, J.S. (2021). Programmable RNA targeting with the single-protein CRISPR effector Cas7-11. Nature 597, 720–725.

6. 6. van Beljouw, S.P.B., Haagsma, A.C., Rodríguez-Molina, A., van den Berg, D.F., Vink, J.N.A., and Brouns, S.J.J. (2021). The gRAMP CRISPR-Cas effector is an RNA endonuclease complexed with a caspase-like peptidase. Science 373, 1349–1353.

7. Pul, Ü., Wurm, R., Arslan, Z., Geissen, R., Hofmann, N., and Wagner, R. (2010). Identification and characterization of *E. coli* CRISPR-cas promoters and their silencing by H-NS. Mol. Microbiol. 75, 1495–1512.

8. Westra, E.R., Pul, Ü., Heidrich, N., Jore, M.M., Lundgren, M., Stratmann, T., Wurm, R., Raine, A., Mescher, M., and Van Heereveld, L. (2010). H-NS-mediated repression of CRISPR-based immunity in *Escherichia coli* K12 can be relieved by the transcription activator LeuO. Mol. Microbiol. 77, 1380–1393.

9. MacRitchie, D.M., Buelow, D.R., Price, N.L., and Raivio, T.L. (2008). Two-component signaling and gram-negative envelope stress response systems. Adv. Exp. Med. Biol. 631, 80–110.

10. Perez-Rodriguez, R., Haitjema, C., Huang, Q., Nam, K.H., Bernardis, S., Ke, A., and DeLisa, M.P. (2011). Envelope stress is a trigger of CRISPR RNA-mediated DNA silencing in *Escherichia coli*. Mol. Microbiol. 79, 584–599.

11. Liu, T., Liu, Z., Ye, Q., Pan, S., Wang, X., Li, Y., Peng, W., Liang, Y., She, Q., and Peng, N. (2017). Coupling transcriptional activation of CRISPR–Cas system and DNA repair genes by Csa3a in *Sulfolobus islandicus*. Nucl.Acids Res. 45, 8978– 8992.

12. He, F., Vestergaard, G., Peng, W., She, Q., and Peng, X. (2017). CRISPR-Cas type I-A Cascade complex couples viral infection surveillance to host transcriptional regulation in the dependence of Csa3b. Nucl. Acids Res. 45, 1902– 1913.

13. Smith, L.M., Hampton, H.G., Yevstigneyeva, M.S., Mahler, M., Paquet, Z.S.M., and Fineran, P.C. (2023). CRISPR-Cas immunity is repressed by the LysR-type transcriptional regulator PigU. Nucl. Acids Res., gkad1165. 10.1093/nar/gkad1165.

14. Patterson, A.G., Chang, J.T., Taylor, C., and Fineran, P.C. (2015). Regulation of the Type I-F CRISPR-Cas system by CRP-cAMP and GalM controls spacer acquisition and interference. Nucl. Acids Res. 43, 6038–6048.

15. Høyland-Kroghsbo, N.M., Paczkowski, J., Mukherjee, S., Broniewski, J., Westra, E., Bondy-Denomy, J., and Bassler, B.L. (2017). Quorum sensing controls the *Pseudomonas aeruginosa* CRISPR-Cas adaptive immune system. PNAS 114, 131–135.

16. Høyland-Kroghsbo, N.M., Muñoz, K.A., and Bassler, B.L. (2018). Temperature, by controlling growth rate, regulates CRISPR-Cas activity in *Pseudomonas aeruginosa*. mBio 9, e02184–18. 1

17. Murray, A.G., and Eldridge, P.M. (1994). Marine viral ecology: incorporation of bacteriophage into the microbial planktonic food web paradigm. J. Plankt. Res. 16, 627–641.

18. Carlson, M.C.G., Ribalet, F., Maidanik, I., Durham, B.P., Hulata, Y., Ferrón, S., Weissenbach, J., Shamir, N., Goldin, S., Baran, N., et al. (2022). Viruses affect picocyanobacterial abundance and biogeography in the North Pacific Ocean. Nat. Microbiol. 7, 570–580.

19. Scholz, I., Lange, S.J., Hein, S., Hess, W.R., and Backofen, R. (2013). CRISPR-Cas systems in the cyanobacterium Synechocystis sp. PCC6803 exhibit distinct processing pathways involving at least two Cas6 and a Cmr2 protein. PLoS ONE 8, e56470.

20. Reimann, V., Alkhnbashi, O.S., Saunders, S.J., Scholz, I., Hein, S., Backofen, R., and Hess, W.R. (2017). Structural constraints and enzymatic promiscuity in the Cas6-dependent generation of crRNAs. Nucl. Acids Res. 45, 915–925.

21. Kieper, S.N., Almendros, C., Behler, J., McKenzie, R.E., Nóbrega, F.L., Haagsma, A.C., Vink, J.N.A., Hess, W.R., and Brouns, S.J.J. (2018). Cas4 facilitates PAM-compatible spacer selection during CRISPR adaptation. Cell Rep. 22, 3377–338.

22. Behler, J., Sharma, K., Reimann, V., Wilde, A., Urlaub, H., and Hess, W.R. (2018). The host-encoded RNase E endonuclease as the crRNA maturation enzyme in a CRISPR–Cas subtype III-Bv system. Nat. Microbiol. 3, 367–377.

23. Scholz, I., Lott, S.C., Behler, J., Gärtner, K., Hagemann, M., and Hess, W.R. (2019). Divergent methylation of CRISPR repeats and cas genes in a subtype I-D CRISPR-Cas-system. BMC Microbiol. 19, 147.1-147.11.

24. McBride, T.M., Schwartz, E.A., Kumar, A., Taylor, D.W., Fineran, P.C., and Fagerlund, R.D. (2020). Diverse CRISPR-Cas complexes require independent translation of small and large subunits from a single gene. Mol. Cell 80, 971–979.

25. Schwartz, E., Bravo, J., Ahsan, M., Macias, L., McCafferty, C., Dangerfield, T., Walker, J., Brodbelt, J., Palermo, G., Fineran, P., et al. (2023). Type III CRISPR-Cas effectors act as protein-assisted ribozymes during RNA cleavage. Res. Sq., rs.3.rs-2837968.

26. Hein, S., Scholz, I., Voß, B., and Hess, W.R. (2013). Adaptation and modification of three CRISPR loci in two closely related cyanobacteria. RNA Biol. 10, 852–864.

27. Makarova, K.S., Anantharaman, V., Grishin, N.V., Koonin, E.V., and Aravind, L. (2014). CARF and WYL domains: ligand-binding regulators of prokaryotic defense systems. Front. Genet. 5, 102.

28. Garcia-Doval, C., Schwede, F., Berk, C., Rostøl, J.T., Niewoehner, O., Tejero, O., Hall, J., Marraffini, L.A., and Jinek, M. (2020). Activation and self-inactivation mechanisms of the cyclic oligoadenylate-dependent CRISPR ribonuclease Csm6. Nat. Commun. 11, 1596.

29. Ding, J., Schuergers, N., Baehre, H., and Wilde, A. (2022). Enzymatic properties of CARF-domain proteins in Synechocystis sp. PCC 6803. Front. Microbiol. 13, 1046388.

30. Riediger, M., Kadowaki, T., Nagayama, R., Georg, J., Hihara, Y., and Hess, W.R. (2019). Biocomputational analyses and experimental validation identify the regulon controlled by the redox-responsive transcription factor RpaB. iScience 15, 316– 331.

31. Kujat, S.L., and Owttrim, G.W. (2000). Redox-regulated RNA helicase expression. Plant Physiol. 124, 703–714.

32. Chamot, D., Colvin, K.R., Kujat-Choy, S.L., and Owttrim, G.W. (2005). RNA structural rearrangement via unwinding and annealing by the cyanobacterial RNA helicase, CrhR. J. Biol. Chem. 280, 2036–2044.

33. Prakash, J.S.S., Krishna, P.S., Sirisha, K., Kanesaki, Y., Suzuki, I., Shivaji, S., and Murata, N. (2010). An RNA helicase, CrhR, regulates the low-temperature-inducible expression of heat-shock genes groES, groEL1 and groEL2 in Synechocystis sp. PCC 6803. Microbiology 156, 442–451.

34. Georg, J., Rosana, A.R.R., Chamot, D., Migur, A., Hess, W.R., and Owttrim, G.W. (2019). Inactivation of the RNA helicase CrhR impacts a specific subset of the transcriptome in the cyanobacterium Synechocystis sp. PCC 6803. RNA Biol. 16, 1205–1214.

35. Rowland, J.G., Simon, W.J., Prakash, J.S.S., and Slabas, A.R. (2011). Proteomics reveals a role for the RNA helicase crhR in the modulation of multiple metabolic pathways during cold acclimation of Synechocystis sp. PCC6803. J. Proteome Res. 10, 3674–3689.

36. Migur, A., Heyl, F., Fuss, J., Srikumar, A., Huettel, B., Steglich, C., Prakash, J.S.S., Reinhardt, R., Backofen, R., Owttrim, G.W., et al. (2021). The temperature-regulated DEAD-box RNA helicase CrhR interactome: Autoregulation and photosynthesis-related transcripts. J Exp Bot 72, 7564–7579.

37. Kopf, M., Klähn, S., Scholz, I., Matthiessen, J.K.F., Hess, W.R., and Voß, B. (2014). Comparative analysis of the primary transcriptome of Synechocystis sp. PCC 6803. DNA Res. 21, 527–539.

38. Kunert, A., Hagemann, M., and Erdmann, N. (2000). Construction of promoter probe vectors for *Synechocystis* sp. PCC 6803 using the light-emitting reporter systems Gfp and LuxAB. J. Microbiol. Methods 41, 185–194.

39. Kadowaki, T., Nagayama, R., Georg, J., Nishiyama, Y., Wilde, A., Hess, W.R., and Hihara, Y. (2016). A feed-forward loop consisting of the response regulator RpaB and the small RNA PsrR1 controls light acclimation of photosystem I gene expression in the cyanobacterium *Synechocystis* sp. PCC 6803. Plant Cell Physiol. 57, 813–823.

40. Eriksson, J., Salih, G.F., Ghebramedhin, H., and Jansson, C. (2000). Deletion mutagenesis of the 5′ *psbA2* region in *Synechocystis* 6803: Identification of a putative cis element involved in photoregulation. Mol. Cell Biol. Res. Comm. 3, 292–298.

41. Song, K., Baumgartner, D., Hagemann, M., Muro-Pastor, A.M., Maaß, S., Becher, D., and Hess, W.R. (2022). AtpΘ is an inhibitor of F0F1 ATP synthase to arrest ATP hydrolysis during low-energy conditions in cyanobacteria. Curr. Biol.32, 136–148.e5.

42. Song, K., Hagemann, M., Georg, J., Maaß, S., Becher, D., and Hess, W.R. (2022). Expression of the cyanobacterial FoF1 ATP synthase regulator AtpΘ depends on small DNA-binding proteins and differential mRNA stability. Microbiol. Spectr. 10, e0256221.

43. Langmead, B., and Salzberg, S.L. (2012). Fast gapped-read alignment with Bowtie 2. Nat. Methods 9, 357–359. 1.

44. Bischler, T. (2022). PEAKachu. https://github.com/tbischler/PEAKachu

45. Rosana, A.R.R., Whitford, D.S., Migur, A., Steglich, C., Kujat-Choy, S.L., Hess, W.R., and Owttrim, G.W. (2020). RNA helicase-regulated processing of the Synechocystis rimO-crhR operon results in differential cistron expression and accumulation of two sRNAs. J. Biol. Chem. 295, 6372–6386.

46. Sharma, K., Hrle, A., Kramer, K., Sachsenberg, T., Staals, R.H.J., Randau, L., Marchfelder, A., van der Oost, J., Kohlbacher, O., Conti, E., et al. (2015). Analysis of protein-RNA interactions in CRISPR proteins and effector complexes by UV-induced cross-linking and mass spectrometry. Methods 89, 138–148.

47. Jumper, J., Evans, R., Pritzel, A., Green, T., Figurnov, M., Ronneberger, O., Tunyasuvunakool, K., Bates, R., Žídek, A., Potapenko, A., et al. (2021). Highly accurate protein structure prediction with AlphaFold. Nature 596, 583–589.

48. Mirdita, M., Schütze, K., Moriwaki, Y., Heo, L., Ovchinnikov, S., and Steinegger, M. (2022). ColabFold: making protein folding accessible to all. Nat. Methods 19, 679–682.

49. Whitman, B.T., Murray, C.R.A., Whitford, D.S., Paul, S.S., Fahlman, R.P., Glover, M.J.N., and Owttrim, G.W. (2022). Degron-mediated proteolysis of CrhR-like DEAD-box RNA helicases in cyanobacteria. J. Biol. Chem. 298, 101925.

50. Huen, J., Lin, C.-L., Golzarroshan, B., Yi, W.-L., Yang, W.-Z., and Yuan, H.S. (2017). Structural Insights into a unique dimeric DEAD-box helicase CshA that promotes RNA decay. Structure 25, 469–481. 1

51. Yang, C.-D., Chen, Y.-H., Huang, H.-Y., Huang, H.-D., and Tseng, C.-P. (2014). CRP represses the CRISPR/Cas system in *Escherichia coli*: evidence that endogenous CRISPR spacers impede phage P1 replication. Mol. Microbiol. 92, 1072–1091. 1

52. Wilde, A., and Hihara, Y. (2016). Transcriptional and posttranscriptional regulation of cyanobacterial photosynthesis. Biochim. Biophys. Acta 1857, 296–308.

53. Lillestøl, R.K., Shah, S.A., Brügger, K., Redder, P., Phan, H., Christiansen, J., and Garrett, R.A. (2009). CRISPR families of the crenarchaeal genus *Sulfolobus*: bidirectional transcription and dynamic properties. Mo. Microbiol. 72, 259–272.

54. Erdmann, S., and Garrett, R.A. (2012). Selective and hyperactive uptake of foreign DNA by adaptive immune systems of an archaeon via two distinct mechanisms. Mol. Microbiol. 85, 1044–1056.

55. Díez-Villaseñor, C., Guzmán, N.M., Almendros, C., García-Martínez, J., and Mojica, F.J.M. (2013). CRISPR-spacer integration reporter plasmids reveal distinct genuine acquisition specificities among CRISPR-Cas I-E variants of *Escherichia coli*. RNA Biol. 10, 792–802.

56. Yosef, I., Shitrit, D., Goren, M.G., Burstein, D., Pupko, T., and Qimron, U. (2013). DNA motifs determining the efficiency of adaptation into the *Escherichia coli* CRISPR array. PNAS 110, 14396–14401.

57. Lin, P., Pu, Q., Wu, Q., Zhou, C., Wang, B., Schettler, J., Wang, Z., Qin, S., Gao, P., Li, R., et al. (2019). High-throughput screen reveals sRNAs regulating crRNA biogenesis by targeting CRISPR leader to repress Rho termination. Nat. Commun. 10, 3728.

58. Kramer, K., Sachsenberg, T., Beckmann, B.M., Qamar, S., Boon, K.-L., Hentze, M.W., Kohlbacher, O., and Urlaub, H. (2014). Photo-cross-linking and high-resolution mass spectrometry for assignment of RNA-binding sites in RNA-binding proteins. Nat. Methods 11, 1064–1070..

59. Linder, P., and Jankowsky, E. (2011). From unwinding to clamping - the DEAD box RNA helicase family. Nat. Rev. Mol. Cell Biol. 12, 505–516.

60. Shchepachev, V., Bresson, S., Spanos, C., Petfalski, E., Fischer, L., Rappsilber, J., and Tollervey, D. (2018). Defining the RNA interactome by total RNA-associated protein purification. Preprint at bioRxiv, 10.1101/436253.

61. Knörlein, A., Sarnowski, C.P., de Vries, T., Stoltz, M., Götze, M., Aebersold, R., Allain, F.H.-T., Leitner, A., and Hall, J. (2022). Nucleotide-amino acid π-stacking interactions initiate photo cross-linking in RNA-protein complexes. Nat. Commun. 13, 2719. 1

62. Bourgeois, C.F., Mortreux, F., and Auboeuf, D. (2016). The multiple functions of RNA helicases as drivers and regulators of gene expression. Nat. Rev. Mol. Cell Biol. 17, 426–438.

63. Khemici, V., and Linder, P. (2018). RNA helicases in RNA decay. Biochem. Soc. Trans. 46, 163–172.

64. Rosana, A.R.R., Chamot, D., and Owttrim, G.W. (2012). Autoregulation of RNA helicase expression in response to temperature stress in *Synechocystis* sp. PCC 6803. PLoS ONE 7, e48683.

65. Sireesha, K., Radharani, B., Krishna, P.S., Sreedhar, N., Subramanyam, R., Mohanty, P., and Prakash, J.S.S. (2012). RNA helicase, CrhR is indispensable for the energy redistribution and the regulation of photosystem stoichiometry at low temperature in *Synechocystis* sp. PCC6803. Biochim. Biophys. Acta 1817, 1525– 1536.

66. Rosana, A.R.R., Whitford, D.S., Fahlman, R.P., and Owttrim, G.W. (2016). Cyanobacterial RNA helicase CrhR localizes to the thylakoid membrane region and cosediments with degradosome and polysome complexes in *Synechocystis* sp. strain PCC 6803. J. Bacteriol. 198, 2089–2099. 10.1128/JB.00267-16.

67. Clokie, M.R.J., and Mann, N.H. (2006). Marine cyanophages and light. Environmental Microbiology 8, 2074–2082.

68. Chou-Zheng, L., and Hatoum-Aslan, A. (2019). A type III-A CRISPR-Cas system employs degradosome nucleases to ensure robust immunity. Elife 8, e45393.

69. Stanier, R.Y., Deruelles, J., Rippka, R., Herdman, M., and Waterbury, J.B. (1979). Generic assignments, strain histories and properties of pure cultures of cyanobacteria. Microbiology 111, 1–61.

70. Zhang, L., McSpadden, B., Pakrasi, H.B., and Whitmarsh, J. (1992). Copper-mediated regulation of cytochrome c553 and plastocyanin in the cyanobacterium *Synechocystis* 6803. J. Biol. Chem. 267, 19054–19059.

71. Pinto, F., Thapper, A., Sontheim, W., and Lindblad, P. (2009). Analysis of current and alternative phenol based RNA extraction methodologies for cyanobacteria. BMC Mol. Biol. 10, 79. 1

72. Kelly, C.L., Taylor, G.M., Hitchcock, A., Torres-Méndez, A., and Heap, J.T. (2018). A rhamnose-inducible system for precise and temporal control of gene expression in cyanobacteria. ACS Synth. Biol. 7, 1056–1066.

73. Beyer, H.M., Gonschorek, P., Samodelov, S.L., Meier, M., Weber, W., and Zurbriggen, M.D. (2015). AQUA cloning: a versatile and simple enzyme-free cloning approach. PLOS ONE 10, e0137652.

74. Klähn, S., Baumgartner, D., Pfreundt, U., Voigt, K., Schön, V., Steglich, C., and Hess, W.R. (2014). Alkane biosynthesis genes in cyanobacteria and their transcriptional organization. Front. Bioeng. Biotechnol. 2, 24.

75. Martin, M. (2011). Cutadapt removes adapter sequences from high-throughput sequencing reads. EMBnet j. 17.

76. Zhan, J., Steglich, C., Scholz, I., Hess, W.R., and Kirilovsky, D. (2021). Inverse regulation of light harvesting and photoprotection is mediated by a 3’-end-derived sRNA in cyanobacteria. Plant Cell 33, 358–380.

77. Bäumert, H.G., Sköld, S.-E., and Kurland, C.G. (1978). RNA-protein neighbourhoods of the ribosome obtained by crosslinking. Eur. J. Biochem. 89, 353–359.

