## Supplementary Information for "Control of a type III-Dv CRISPR‒Cas system by the transcription factor RpaB and interaction of its leader transcript with the DEAD-box RNA helicase CrhR"

### Supplementary Tables

**Table S1.** *In vitro* RNA-CrhR pulldown. Absolute and relative numbers of reads derived from RNA-seq data analysis.

| | | **Mapped to *Synechocystis*** | | **Mapped to *E. coli*** | |
|---|---|---|---|---|---|
| **Sample** | Total number | Number of mapped | % of mapped | Number of mapped | % of mapped |
| **CrhR 1** | 10,753,338 | 697,535 | 6.5 | 9,742,457 | 90.6 |
| **CrhR 2** | 10,742,452 | 1,242,204 | 11.6 | 9,175,217 | 85.4 |
| **CrhR$_{K57A}$ 1** | 10,575,705 | 5,378,287 | 50.9 | 4,905,923 | 46.4 |
| **CrhR$_{K57A}$ 2** | 10,725,162 | 5,187,642 | 48.4 | 5,272,250 | 49.2 |
| ***Synechocystis* total RNA 1** | 10,739,872 | 10,174,598 | 94.7 | 203,419 | 1.9 |
| ***Synechocystis* total RNA 2** | 10,741,620 | 10,232,016 | 95.3 | 285,748 | 2.7 |

**Table S2.** Oligonucleotide primers used in this work.

| Number | Name | Sequence |
|---|---|---|
| PCR primers for generation of expression vectors | | |
| 1 | 3xFL-CrhR-F1 | GATTACAAGGATGACGATGACAAGACTAATACTTTGA CTAGTACCTTCG |
| 2 | 3xFL-CrhR-F2 | GATTATAAAGATCATGATATCGATTACAAGGATGACG ATGACAAGAC |
| 3 | 3xFL-CrhR-F3 | GCCA↓TATGGACTACAAAGACCATGACGGTGATTATA AAGATCATGATATC (*Nde*I*) |
| 4 | 3xFL-CrhR-R | CCGT↓CTAGACCAGAGCT↓CATTACTGTTGGCGATCA CTATAGGCAGGACG (*Xba*I, *Sac*I*) |
| 5 | crhR$_{K57A}$_inverse_fw | AATGCGGTCCATCAAAGGTAGGGCAAAGGCGGCGG TTGCCCCCGTGCCGGTTTGGG |
| 6 | crhR$_{K57A}$_inverse_rv | GACCCAGAGGGGGACTTGCAAGCCTTAATTTTGACC |
| 7 | petE-3xF-rrnb_neg_ctrl_FP | GCCGAGCT↓CGCCTGGCGGCAG (*Sac*I*) |
| 8 | petE-3xF-rrnb_neg_ctrl_RP | GGCGAGCT↓CTTACTTGTCATCGTCATCCTTGTAATC GATATCA (*Sac*I*) |
| 9 | petE_foR-pVZ_FP | CGGG↓AATTCCAAGGCAAAACCACCGTTATCAGCAG (*Eco*RI*) |
| 10 | rrnB_foR-pVZ_RP | CGGG↓AATTCAAAAAGGCCATCCGTCAGGATG (*Eco*RI*) |
| 11 | pQE70_crhR-fw | GAAATTAAGCATGACTAATACTTTGACTAGTACC |
| 12 | pQE70_crhR-rv | TGGTGATGAGATCTCTGTTGGCGATCACTATAGG |
| 13 | pQE_GIBSOn-fw | AGATCTCATCACCATCACCATC |
| 14 | pQE_Gibson-rv | GCTTAATTTCTCCTCTTTAATGAATTCTGTG |
| PCR primers and oligonucleotides to generate riboprobes for northern blots | | |
| 15 | crhR_NB_Fw | TAATACGACTCACTATAGGGGGTCAAAATTAAGGCTT GCAAGT |
| 16 | crhR_NB_Rv | GAAGCCATTCCCCTGCTTTTGAGC |
| 17 | ncr0700_northern_for | TAATACGACTCACTATAGGGGACCGGAGCTGCAAAT ACCTAATAC |
| 18 | ncr0700_northern_rev | CTAAATCTGTCTCGTTTGTTTCTCAGTTC |
| 19 | ncl1330_NB_Fw | TAATACGACTCACTATAGGGCTAAATAGGTCAAATTG CAGCG |
| 20 | ncl1330_NB_Rv | CTAAGCTATCTAAGGACCAACACGAC |

| | | |
|---|---|---|
| 21 | syr11_Fw_NB | TAATACGACTCACTATAGGcaaaatgacctgatccccgtgtttttattatgttaagggtctttataaaggtgaccccgtgttatttcgtt |
| 22 | syr11_Rv_NB | AACGAAATAACACGGGGTCACCTTTATAAAGACCCTTAACATAATAAAAAACACGGGGATCAGGTCATTTTGCCTATAGTGAGTCGTATTA |
| 23 | CRISPRlead_NB_Fw | TAATACGACTCACTATAGGGGAAAGTGGAGGAAAGCAGAC |
| 24 | CRISPRlead_NB_Rv | GGTTGCCAAATTGCTTCTGAACCTTG |
| PCR primers and oligonucleotides to create T7 transcription DNA template or DNA target for EMSA | | |
| 25 | RNaseE_CrhR_T7_Fw | TAATACGACTCACTATAGGTATGACCGCCGAGGAGGCTAAGGTTTTTTAGCTATTGAA |
| 26 | RNaseE_CrhR_T7_Rv | TTCAATAGCTAAAAAACCTTAGCCTCCTCGGCGGTCATACCTATAGTGAGTCGTATTA |
| 27 | EMSA_CrhR-T7_Fw | TAATACGACTCACTATAGGATGACTAATACTTTGACTAGTACCTTC |
| 28 | EMSA_CrhR-T7_Rv | TTGGGCCAGCATGTCCCG |
| 29 | EMSA_CRISPR2LeadeR-T7_Fw | TAATACGACTCACTATAGGTATAGCCAAAAATAAGGCTTTTTATTGTTATG |
| 30 | EMSA_CRISPR2LeadeR-T7_Rv | CCTATCACACTGAAAAACAGTCTG |
| 31 | EMSA_CRISPR2spaceR-T7_Fw | TAATACGACTCACTATAGGACTGAAACCTTGGTATTTGTAGTTCTCGATGAGTGTTTTAGGCAGTTCAACACCCTC |
| 32 | EMSA_CRISPR2spaceR-T7_Rv | GAGGGTGTTGAACTGCCTAAAACACTCATCGAGAACTACAAATACCAAGGTTTCAGTCCTATAGTGAGTCGTATTA |
| 33 | EMSA_P*cas10*_HLR1_fw | GTATCCCCACCCCTGTTGATTAAAC |
| 34 | EMSA_P*cas10*_HLR1_rev | TTCCGAAAGGGGATCAATTCTTGG |
| 35 | EMSA_P*psbA2*_fw | CATCTATAAGCTTCGTGTATATTAAC |
| 36 | EMSA_P*psbA2*_rev | ACTAAACTTAGTCTAAAGGATTAATG |
| 37 | CRISPR2LeadeR-helix_T7_Fw | TAATACGACTCACTATAGGTTTTTATTGTTATGTTTTCAGTACGTAACAAATC |
| 38 | CRISPR2LeadeR-helix_T7_Rv | GATTTGTTACGTACTGAAAACATAACAATAAAAACCTATAGTGAGTCGTATTA |
| 39 | CRISPR2LeadeR-dishelix_T7_Fw | TAATACGACTCACTATAGGTTTTTATTGTTATGTTTTCAGTACTGAACAAATC |
| 40 | CRISPR2LeadeR-dishelix_T7_Rv | GATTTGTTCAGTACTGAAAACATAACAATAAAAACCTATAGTGAGTCGTATTA |
| 41 | CRISPR2LeadeR-inline_Fw | TAATACGACTCACTATAGGTATAGCCAAAAATAAGGCTTTTTATTGTTATGTTTTCAGTACGTAACAAATCGTTAAACGTCTACCTCTTGTTTGGCGGATTGGCACCTCGAAAACCGCA |

| | | |
|---|---|---|
| 42 | CRISPR2LeadeR-inline_Rv | TGCGGTTTTCGAGGTGCCAATCCGCCAAACAAGAGGTAGACGTTTAACGATTTGTTACGTACTGAAAACATAACAATAAAAAGCCTTATTTTTGGCTATACCTATAGTGAGTCGTATTA |
| DNA/RNA oligonucleotides and PCR primers used to prepare DNA libraries for RNA-seq | | |
| 43 | Loop Primer 7N | AGATCGGAAGAGAGACGTGTGCTCTTCCGATCTNNNNNNN* |
| 44 | Loop primer 6N | AGATCGGAAGAGAGACGTGTGCTCTTCCGATCTNNNNNND* |
| 45 | Loop primer 5N | AGATCGGAAGAGAGACGTGTGCTCTTCCGATCTNNNNNDD* |
| 46 | Loop primer 4N | AGATCGGAAGAGAGACGTGTGCTCTTCCGATCTNNNNDDD* |
| 47 | 5'-adapter | CCCUACACGACGCUCUUCCGAUCUGA (RNA) |
| 48 | SMART oligo | d(CCCTACACGACGCTCTTCCGATCT)-r(GGG) (DNA-RNA fusion) |
| 49 | PCR Primer 1 | AATGATACGGCGACCACCGAGATCTACACTCTTTCCCTACACGACGCTCTTCCGATCTGA |
| 50 | PCR Primer 2.1:R701: ATCACG | CAAGCAGAAGACGGCATACGAGATcgtgatGTGACTGGAGTTCAGACGTGTGCTCTTCCGATCT |
| 51 | PCR Primer 2.2:R702: CGATGT | CAAGCAGAAGACGGCATACGAGATacatcgGTGACTGGAGTTCAGACGTGTGCTCTTCCGATCT |
| 52 | PCR Primer 2.3:R703: TTAGGC | CAAGCAGAAGACGGCATACGAGATgcctaaGTGACTGGAGTTCAGACGTGTGCTCTTCCGATCT |
| 53 | PCR Primer 2.4:R704: TGACCA | CAAGCAGAAGACGGCATACGAGATtggtcaGTGACTGGAGTTCAGACGTGTGCTCTTCCGATCT |
| 54 | PCR Primer 2.5:R705: ACAGTG | CAAGCAGAAGACGGCATACGAGATcactgtGTGACTGGAGTTCAGACGTGTGCTCTTCCGATCT |
| 55 | PCR Primer 2.6:R706: GCCAAT | CAAGCAGAAGACGGCATACGAGATattggcGTGACTGGAGTTCAGACGTGTGCTCTTCCGATCT |
| 56 | PCR Primer 2.7:R707: CAGATC | CAAGCAGAAGACGGCATACGAGATgatctgGTGACTGGAGTTCAGACGTGTGCTCTTCCGATCT |
| 57 | PCR Primer 2.8:R708: ACTTGA | CAAGCAGAAGACGGCATACGAGATtcaagtGTGACTGGAGTTCAGACGTGTGCTCTTCCGATCT |
| 58 | PCR Primer 2.9:R709: GATCAG | CAAGCAGAAGACGGCATACGAGATctgatcGTGACTGGAGTTCAGACGTGTGCTCTTCCGATCT |
| 59 | PCR Primer 2.10:R710: TAGCTT | CAAGCAGAAGACGGCATACGAGATaagctaGTGACTGGAGTTCAGACGTGTGCTCTTCCGATCT |
| Oligonucleotides for cloning pILA vectors | | |
| 60 | pILA_1_fw | AAGTCAAATTTTTATGAAGTTTGGAAATATTTGTTTTTC |
| 61 | pILA_1_rev | TATGGTAAATTCATTTCGATTTTTTGGTTCAAC |
| 62 | pILA_2_fw | AAAAATCGAAATGAATTTACCATAATAAAATTAAAGGC |

| | | |
|---|---|---|
| 63 | pILA_2_rev | ATCCTTTTTTTCTGCGCGTAATCTGCTGCTTGCAAACAAAAAAAC |
| 64 | pILA_3_fw | CAGATTACGCGCAGAAAAAAAGGATCTC |
| 65 | pILA_3_rev | CGACCTGCACTGCCAATTGGATCCCGGGCCCCTGCAGGTCGACTCTAG |
| 66 | prom_cas10_luxAB_fw | CCAATTGGCAGTGCAGGTCGACTCTGTATCCCCACCCCTGTTGATTAAACGATAGTTAACAAATATTTA |
| 67 | prom_cas10_luxAB_rev | TTCCAAACTTCATAAAAATTTGACTTAACGAAAGGGTCGGATAGGACGAAGT |
| 68 | prom_cas10_mut_luxAB_fw | CCAATTGGCAGTGCAGGTCGACTCTGTATCCCCACCCCTGTTGATTAAACGATAGTTAGGGGATATTTA |
| 69 | Prom_CRISPR2_fw | GGGATCCAATTGGCAGTGCAGGTCGGACGTTGTAGCAGTGAGC |
| 70 | Prom_CRISPR2_RBS_rev | TTCTACCTCCTTTGTATATTATAAACTTACCCGAAAAGGTCATAGTAATACCTAAATTTG |
| 71 | pILA_1_RBS_fw | ATAATATACAAAGGAGGTAGAAATGAAGTTTGGAAATATTTGTTTTTCG |
| 72 | pIGA-fw | CTCAGCGCCAAGAGTAGTTCC |
| 73 | pIGA-rev | CACTCTGCACTGTGTCTGTGC |

### Supplementary Figures

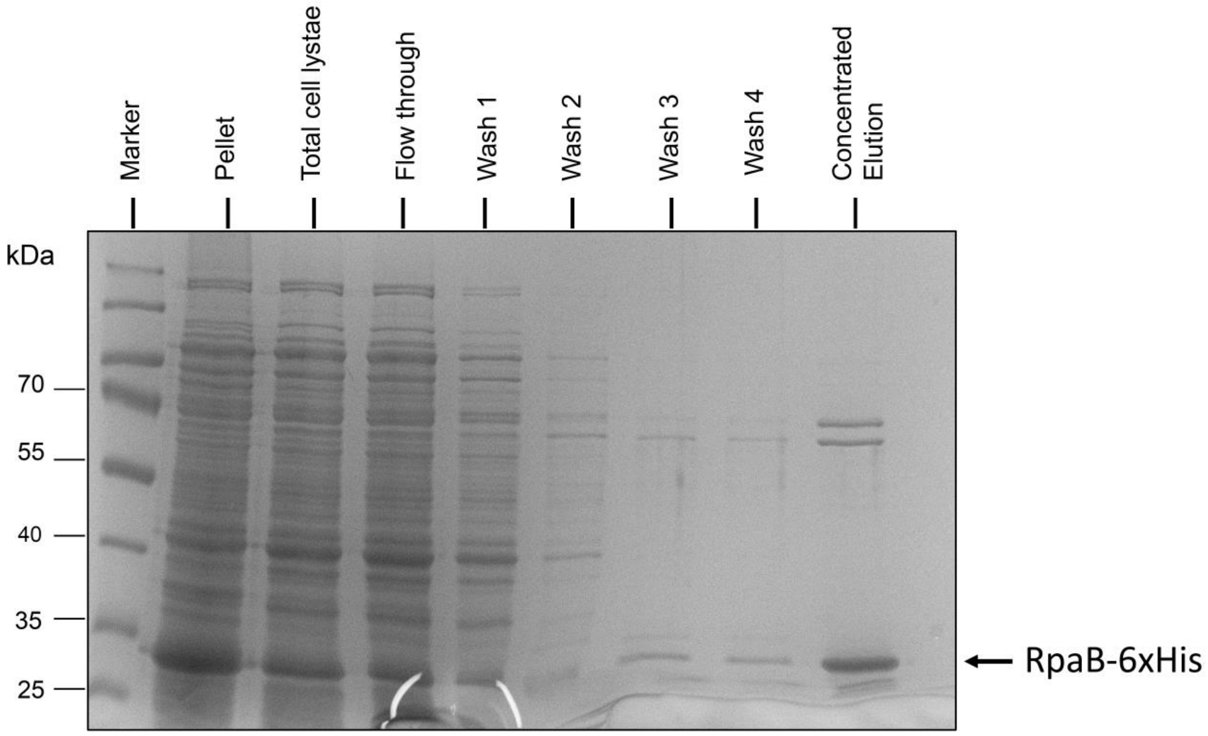


**Figure S1. Expression and purification of recombinant RpaB.** 6×His-RpaB from *Synechocystis* 6803 was expressed and purified from Rosetta (DE3) + pLysS *E. coli* strain. Overexpression was induced with 1 mM IPTG when culture reached $OD_{600}$ 0.6 and continued cultivation for 3 hours at 37°C. Histidine tagged proteins were purified via affinity chromatography using Ni-NTA agarose beads from Qiagen. Column was washed with 40 mM imidazole and target proteins were eluted with 300 mM imidazole. Samples from soluble crude extract, flow through, wash and elution fractions were analysed on denaturing 15% polyacrylamide gels and stained with Instant Coomassie Protein Stain. M: PageRuler Prestained Protein Ladder, P: pellet, sCE: soluble crude extract, FT: Flow through, W: wash and E: elution. We assessed the relative amount of RpaB in the elution fractions, as other proteins were eluted as well. Using Image J[1], we estimated the purity of RpaB in the elution fraction to be approximately 72%.

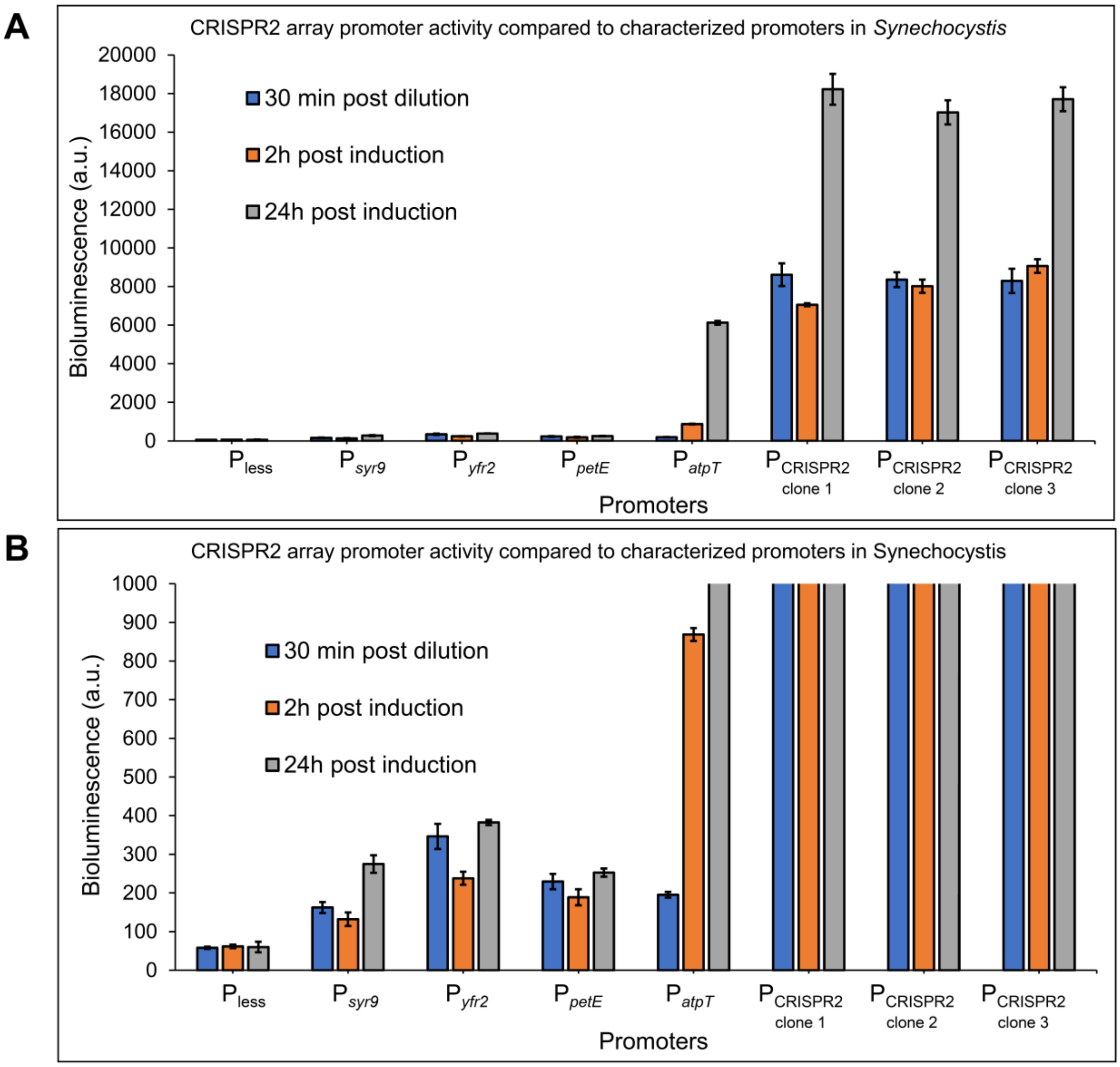


**Figure S2. Comparative *luxAB* promoter assays with the CRISPR2 array promoter and promoters characterized previously.** Cultures were diluted to $OD_{750nm}$ = 0.4 and incubated for 30 min under low light before measuring basal luminescence (30 min post dilution). Cells were then induced according to their stimulator: 2 µmol of copper for $P_{petE}$, cultures with $P_{atpT}$ were wrapped in aluminium foil (darkness). For $P_{syr9}$, $P_{yfr2}$ and $P_{CRISPR2}$, cultures were kept in standard growth conditions as previous data showed that their respective transcripts accumulate the most during the exponential phase[2]. Bioluminescence was measured again after 2 hours post induction and 24 hours post induction. **A.** Graphical representation of the results. **B.** Zoom in into the same data as in panel A to visualize the activity of the promoters $P_{syr9}$, $P_{yfr2}$, $P_{petE}$ and $P_{atpT}$, compared to the negative control, $P_{less}$.

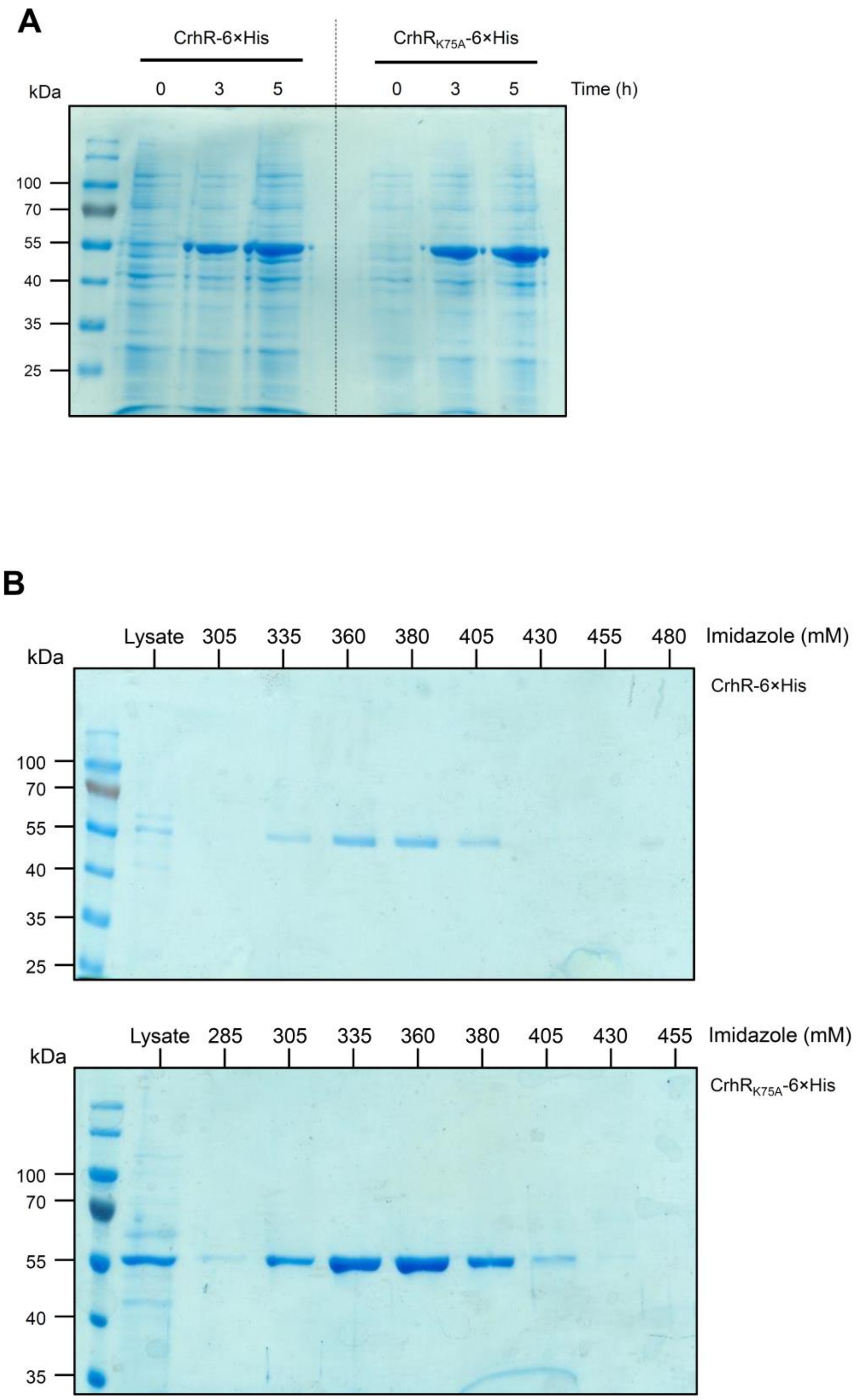


**Figure S3. Expression and purification of recombinant CrhR-6xHis and CrhR$_{K57A}$-6xHis in *E. coli* M15. A.** Soluble proteins isolated from the cell lysate (10 µg) were separated on a 10% SDS polyacrylamide gel and subjected to colloidal Coomassie G-250 staining. Aliquots of the cell cultures were harvested for protein isolation before the addition of IPTG (0 hour) and after 3, and 5 hours of induction with 1 mM IPTG. ~55 kDa proteins corresponding to CrhR were detected in the induced cells. **B.** Purification of recombinant His-CrhR and His-$CrhR_{K57A}$ from *E. coli* M15. *E. coli* cell lysates after 3 hours of IPTG induction and elution fractions obtained with the imidazole concentrations indicated on top are shown. The proteins were separated by SDS-PAGE on 10% polyacrylamide gels and stained with Coomassie G-250.

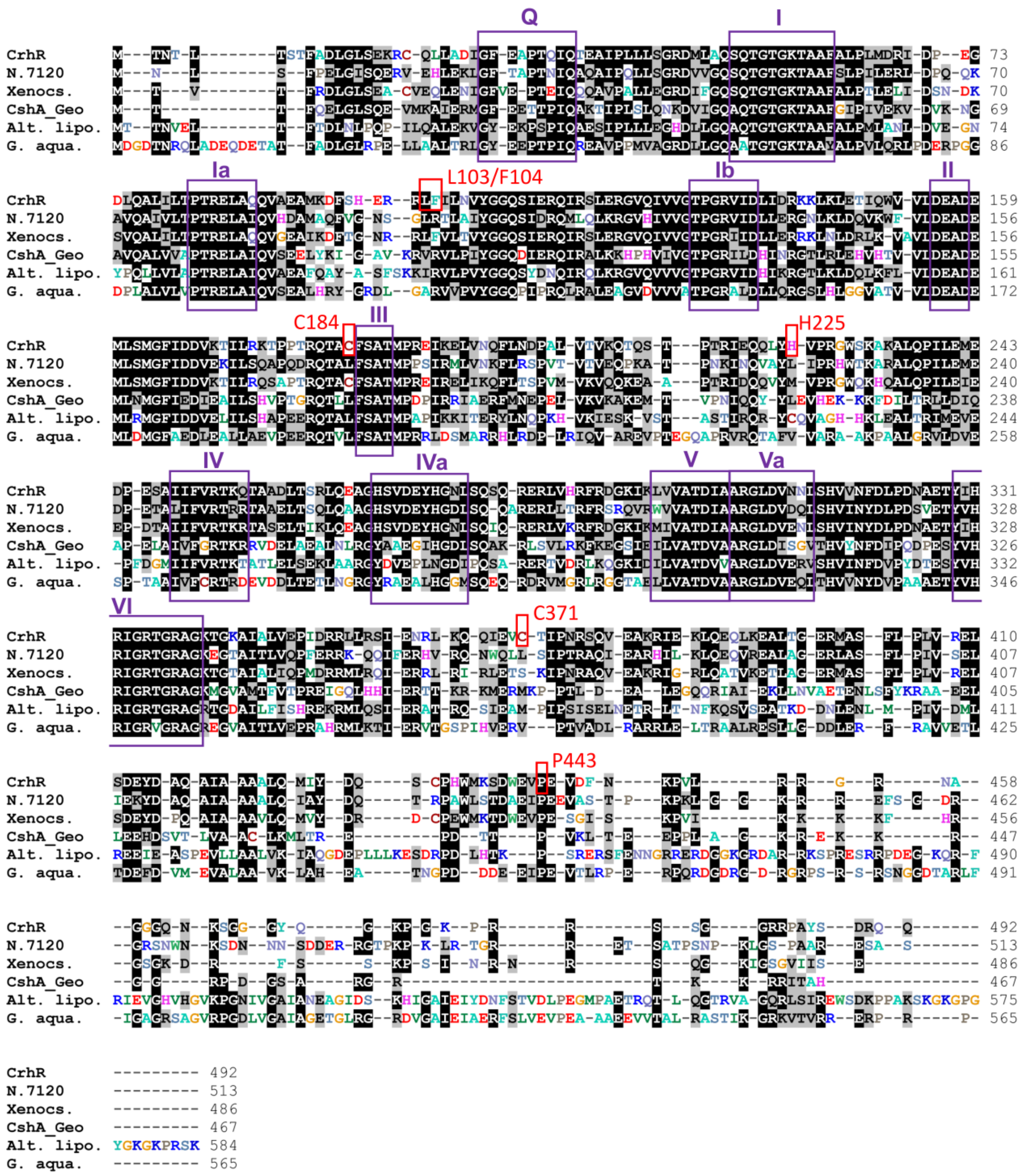


**Figure S4. Cross-linked CrhR residues in the context of previously identified functionally relevant domains of DEAD-box RNA helicases and sequence conservation.** Five DEAD-box RNA helicases were selected for comparison, from the two cyanobacteria *Nostoc* sp. PCC 7120 (N.7120, accession number WP_010995395) and *Xenococcus* sp. MO_188.B8 (Xenocs., MDJ0897852), from *Geobacillus stearothermophilus* (CshA_Geo, which is structurally characterized[3]), from *Alteromonas lipolytica* (Alt. lipo., WP_070176363) and *Geodermatophilus aquaeductus* (G. aqua., WP_246065722). Conserved DEAD-box RNA helicase domains are highlighted by purple boxes and numbered as previously described[4]. Cross-linked residues found in this study are boxed in red and numbered as in **Figure 7**.

### Supplementary References


1. Schneider, T.D., and Stephens, R.M. (1990). Sequence logos: a new way to display consensus sequences. Nucleic Acids Res. *18*, 6097–6100.

2. Kopf, M., Klähn, S., Scholz, I., Matthiessen, J.K.F., Hess, W.R., and Voß, B. (2014). Comparative analysis of the primary transcriptome of *Synechocystis* sp. PCC 6803. DNA Res. *21*, 527–539.

3. Huen, J., Lin, C.-L., Golzarroshan, B., Yi, W.-L., Yang, W.-Z., and Yuan, H.S. (2017). Structural Insights into a unique dimeric DEAD-box helicase CshA that promotes RNA decay. Struct. *25*, 469–481. 10.1016/j.str.2017.01.012.

4. Linder, P., and Jankowsky, E. (2011). From unwinding to clamping - the DEAD box RNA helicase family. Nat. Rev. Mol. Cell Biol. *12*, 505–516.
